# SARS-CoV-2-specific T cells exhibit phenotypic features reflecting robust helper function, lack of terminal differentiation, and high proliferative potential

**DOI:** 10.1101/2020.06.08.138826

**Authors:** Jason Neidleman, Xiaoyu Luo, Julie Frouard, Guorui Xie, Gurjot Gill, Ellen S. Stein, Matthew McGregor, Tongcui Ma, Ashley F. George, Astrid Kosters, Warner C. Greene, Joshua Vasquez, Eliver Ghosn, Sulggi Lee, Nadia R. Roan

**Author notes:** Co-Correspondence. Equal contribution (Jason Neidleman, Xiaoyu Luo, Julie Frouard, Guorui Xie).

## Abstract

Convalescing COVID-19 patients mount robust T cell responses against SARS-CoV-2, suggesting an important role for T cells in viral clearance. To date, the phenotypes of SARS-CoV-2-specific T cells remain poorly defined. Using 38-parameter CyTOF, we phenotyped longitudinal specimens of SARS-CoV-2-specific CD4+ and CD8+ T cells from nine individuals who recovered from mild COVID-19. SARS-CoV-2-specific CD4+ T cells were exclusively Th1 cells, and predominantly Tcm with phenotypic features of robust helper function. SARS-CoV-2-specific CD8+ T cells were predominantly Temra cells in a state of less terminal differentiation than most Temra cells. Subsets of SARS-CoV-2-specific T cells express CD127, can homeostatically proliferate, and can persist for over two months. Our results suggest that long-lived and robust T cell immunity is generated following natural SARS-CoV-2 infection, and support an important role for SARS-CoV-2-specific T cells in host control of COVID-19.

## INTRODUCTION

The first cases of COVID-19 were reported in December of 2019 in Wuhan, China, and soon thereafter its causative agent was identified as SARS-CoV-2, a beta-coronavirus with 79% sequence identity to the SARS-CoV that had emerged in 2003 (Lu et al., 2020). SARS-CoV-2 has proven to be much more transmissible than its SARS-CoV counterpart, quickly spreading around the world and causing what was declared a pandemic on March 11^th^, 2020. By June of 2020, confirmed cases of COVID-19 surpassed 6 million, nearly 6% of which were fatal (Johns Hopkins Coronavirus Resource Center). The outcome from exposure to SARS-CoV-2 can range from asymptomatic infection to death, most often caused by respiratory failure from acute respiratory distress syndrome (ARDS). Limited genetic diversity has been observed in circulating SARS-CoV-2 strains and genetic variations do not seem to correlate with disease severity (Zhang et al., 2020), suggesting that the variable COVID-19 outcomes are driven by variable host responses.

Many individuals exposed to SARS-CoV-2 are asymptomatic or exhibit only a mild course of disease, suggesting that natural immunity can effectively combat this virus. While most studies on SARS-CoV-2 immunity have focused on the humoral immune response, emerging data suggest T cell-mediated immunity is also likely to play an important role in eliminating the virus. Lymphopenia, as characterized by reduced numbers of CD4+ and CD8+ T cells, is predictive of disease severity (Zhang et al., 2020). In addition, levels of activated T cells increase at the time of SARS-CoV-2 clearance (Thevarajan et al., 2020). Furthermore, the clonality of T cell receptor (TCR) sequences (Huang et al., 2020) is higher in patients with mild rather than severe COVID-19, suggesting a role for antigen-specific T cell responses in symptom resolution. A beneficial role for T cells in combating COVID-19 would be in line with observations that both CD4+ and CD8+ T cells are protective against the closely-related SARS-CoV (Li et al., 2008; Roberts et al., 2007; Zhao et al., 2010). However, the nature of the response is also important, as Th1 responses appear to be protective against SARS-CoV while Th2 responses were associated with immunopathology (Deming et al., 2006; Janice Oh et al., 2012; Yasui et al., 2008). Th17 responses have also been implicated in immunopathology during coronavirus infections (Hotez et al., 2020).

Only a limited number of studies have characterized the SARS-CoV-2-specific T cell response. These studies have focused on the breadth of the T cell response, and detected T cells recognizing spike and non-spike epitopes (Braun et al., 2020; Grifoni et al., 2020; Ni et al., 2020; Peng et al., 2020; Weiskopf et al., 2020). In some (Braun et al., 2020; Grifoni et al., 2020) but not other (Peng et al., 2020) studies, responses were also detected in individuals who had not been infected with SARS-CoV-2, presumably reflecting recognition of cross-reactive epitopes from other coronaviruses. These responses primarily involved non-spike ORFs (Grifoni et al., 2020). ELISA revealed that peptide-treated PBMCs upregulated IFNγ but not IL4 or IL17, suggesting a Th1 response (Grifoni et al., 2020; Weiskopf et al., 2020). Limited phenotyping based on CD45RA and CCR7 suggested SARS-CoV-2-specific CD4+ T cells to be more of the T central memory (Tcm) phenotype, while SARS-CoV-2-specific CD8+ T cells were biased towards terminally-differentiated effector cells (Temra) (Weiskopf et al., 2020). These results are consistent with longitudinal analysis of two COVID-19 individuals by bulk TCR sequencing where clonal TCR sequences (assumed to be SARS-CoV-2 specific) were prevalent among Tcm for CD4+ T cells, and Temra for CD8+ T cells (Minervina et al., 2020). However, the phenotypic features of SARS-CoV-2-specific CD4+ and CD8+ T cells have not been systematically investigated, and therefore little is actually known about the functional properties of these cells and their ability to persist long-term in convalescent individuals.

In this study, we conducted an in-depth phenotypic analysis of SARS-CoV-2-specific CD4+ and CD8+ T cells circulating in the bloodstream of individuals who had recently recovered from COVID-19. This was achieved by combining detection of SARS-CoV-2-specific T cells together with CyTOF, a mass spectrometry-based single-cell phenotyping method that uses antibodies conjugated to metal lanthanides to quantify expression levels of both surface and intracellular proteins (Bendall et al., 2011). As spectral overlap is not a limitation with CyTOF, large phenotyping panels of nearly 40 parameters can be implemented, allowing for a high-resolution view of immune cells and the use of high-dimensional analytical methods to monitor cellular remodeling (Cavrois et al., 2017; Ma et al., 2020b). We report here that SARS-CoV-2-specific CD4+ and CD8+ T cells from convalescent individuals are diverse, exhibit features different from antigen-specific T cells against CMV, include cells with both lymphoid and tissue homing potential, harbor phenotypic features of functional effector cells, and are long-lived and capable of homeostatic proliferation.

## RESULTS

### SARS-CoV-2-specific T cells from convalescent individuals produce IFNγ

Nine convalescent and three uninfected participants (Table S1) were recruited from the UCSF acute COVID-19 Host Immune Response Pathogenesis (CHIRP) study for blood donation. Blood specimens were obtained 20-47 days after the participants tested positive by RT-PCR for SARS-CoV-2. PBMCs were purified from the freshly-isolated blood specimens, and then immediately phenotyped by CyTOF, or stimulated for 6 hours with overlapping peptides in the presence of co-stimulation to enable detection of antigen-specific T cells. Overlapping 15-mer peptides from spike, an immunodominant SARS-CoV-2 antigen (Braun et al., 2020; Grifoni et al., 2020), were used for characterization of the SARS-CoV-2-specific response, while overlapping peptides against pp65, an immunodominant CMV antigen (Sylwester et al., 2005), were used for comparison. To enable high-parameter phenotyping of antigen-specific cells, we modified a recently developed human T cell CyTOF panel (Ma et al., 2020b) so it would detect cells producing the cytokines IFNγ, IL4, or IL17 (Table S2). CD4+ and CD8+ T cells were identified by sequential gating on live, singlet CD3+ cells expressing the corresponding co-receptor (Fig. S1). Spike-specific CD4+ and CD8+ T cells producing IFNγ were detected in convalescent but not uninfected individuals (Fig. 1, Table S3), suggesting a robust spike-specific Th1 response. In contrast, spike-specific CD4+ T cells did not include cells of the Th2 and Th17 lineages, as no IL4-or IL17-producing cells were detected following spike peptide stimulation, although such cells were detected following PMA/ionomycin stimulation (Fig. S2).

**Figure 1.**
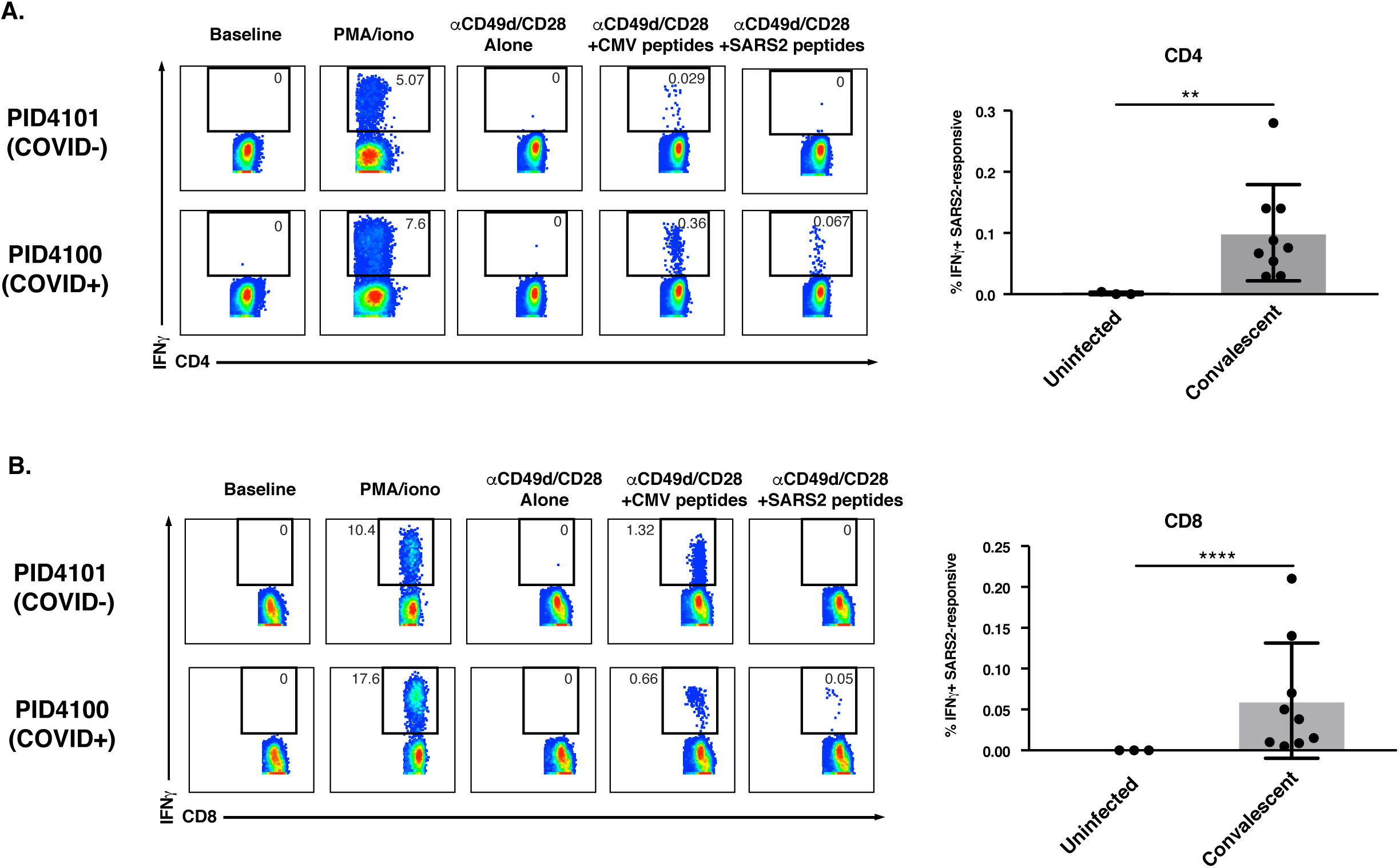
Antigen-specific CD4+ and CD8+ T cells against SARS-CoV-2 spike secrete IFNγ. Shown on the left are pseudocolor plots of CyTOF datasets reflecting the percentage of CD4+ (**A**) or CD8+ (**B**) T cells producing IFNγ in response to the indicated treatment condition, for one representative uninfected (COVID-) and recovered convalescent (COVID+) donor. Numbers correspond to the percentage of cells within the gates. The Baseline condition corresponds to cells phenotyped by CyTOF immediately following isolation of PBMCs from freshly drawn blood, while for all other treatment conditions cells were cultured for 6 hours prior to phenotyping by CyTOF. PMA/ionomycin treatment was used as a positive control. Anti-CD49d/CD28 was used to provide co-stimulation during peptide treatment. Shown on the right are cumulative data from three uninfected individuals and nine recovered convalescent individuals (Table S1). Results are gated on live, singlet CD4+ or CD8+ T cells. ** p < 0.01, **** p < 0.0001 as assessed using the Student’s unpaired t test.

### SARS-CoV-2-specific T cells are phenotypically diverse and different from CMV-specific T cells

To obtain a global view of the phenotypic features of SARS-CoV-2-specific T cells, we visualized the data by t-distributed stochastic neighbor embedding (t-SNE) (van der Maaten and Hinton, 2008). We gated on the IFNγ-producing spike-specific cells (Fig. 1) and overlaid these on total T cells treated with co-stimulation alone. As most of the convalescent donors were CMV+ (Table S1, Table S3), we compared the locations of the spike-specific cells and CMV-specific cells in three representative donors. Both spike-specific CD4+ (Fig. 2A, S3A) and CD8+ (Fig. 2B, S3B) T cells were diverse in that they occupied multiple regions of the t-SNE. However, these cells were concentrated within one or two major regions of the t-SNE, suggesting that their phenotypes are biased towards particular subsets. The phenotypes of spike-specific T cells were not identical to those of CMV-specific T cells, as in every donor there were regions of the t-SNE occupied by CMV-specific T cells that were devoid of spike-specific cells (Fig. 2, *purple ovals*), although overall the CD8+ T cells against the two viruses were more similar to each another than as for the CD4+ T cells. This was further confirmed by demonstrating that the distributions of subset clusters, as defined using the clustering algorithm DensVM (Becher et al., 2014), were different among T cells specific for SARS-CoV-2 as compared to those specific for CMV, particularly for CD4+ T cells (Fig. S4). These results suggest that spike-specific T cells are not randomly distributed among T cell subsets and differ from CMV-specific T cells, a finding that is perhaps expected as CMV differs from SARS-CoV-2. To assess whether the spike-specific T cell response is representative of the overall SARS-CoV-2-specific T cell response, we stimulated in parallel T cells from two convalescent individuals with either the spike peptides, or with a mix of overlapping 15-mer peptides against envelope (env) and nucleocapsid (NC), two other structural proteins of SARS-CoV-2. These results revealed the phenotypes of env/NC-specific cells to be similar to those of cells recognizing spike (Fig. S5). Hereafter, we focus on the spike-specific cells and refer to them as “SARS-CoV-2-specific”.

**Figure 2.**
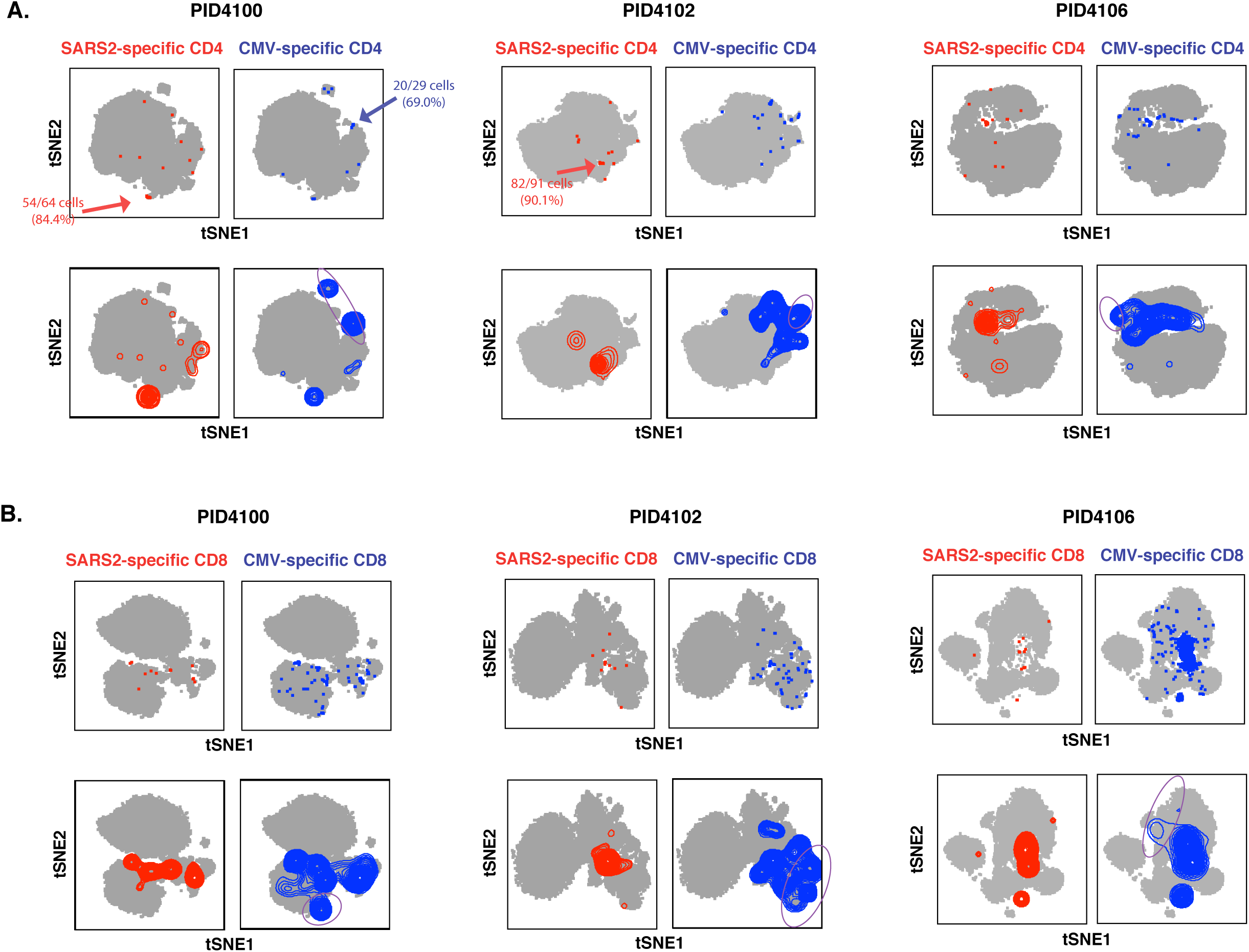
SARS-CoV-2 spike-specific CD4+ and CD8+ T cells are diverse and not phenotypically identical to their CMV-specific counterparts. Shown are t-SNE plots of CyTOF datasets reflecting CD4+ (**A**) or CD8+ (**B**) T cells from three representative COVID-19 convalescent donors who had also sustained a previous CMV infection. Cells shown in grey correspond to CD4+ or CD8+ T cells from specimens stimulated with anti-CD49d/CD28 in the absence of any peptides. The top pairs of plots show SARS-CoV-2 spike-specific (*red*) or CMV pp65-specific (*blue*) cells as individual dots, with some regions concentrated in antigen-specific cells indicated. The bottom pairs of plots show the same data but with the antigen-specific cells shown as contours instead of dots, to better visualize regions with highest densities of antigen-specific cells. Purple ovals outline examples of regions harboring CMV-specific but not SARS-CoV-2-specific cells.

### SARS-CoV-2-specific Th1 cells exhibit phenotypic features characteristic of lymph node homing, robust helper function, and longevity

We then characterized the specific phenotypic features of SARS-CoV-2-specific CD4+ T cells. These cells expressed high levels of the transcription factor Tbet, which together with their ability to induce IFNγ (Fig. 3A) confirms their Th1 differentiation state. These cells, like their CMV-specific counterparts, almost exclusively expressed high levels of CD45RO and low levels of CD45RA, suggesting they are memory cells. Interestingly, while most of the SARS-CoV-2-specific cells expressed CD27 and CCR7, the vast majority of the CMV-specific cells did not (Fig. 3B). This suggests that SARS-CoV-2-specific CD4+ T cells are mostly Tcm, memory cells that home to lymph nodes where they can help B cells undergo affinity maturation. To evaluate the potential helper function of these cells, we assessed their expression of CXCR5 and ICOS, markers of circulating T follicular helper cells (cTfh), cells that efficiently induce viral-specific memory B cells to differentiate into plasma cells and whose levels are associated with protective antibody responses (Bentebibel et al., 2013). SARS-CoV-2-specific CD4+ T cells expressed higher levels of CXCR5 and ICOS than both total and CMV-specific CD4+ T cells (Fig. 3C).

**Figure 3.**
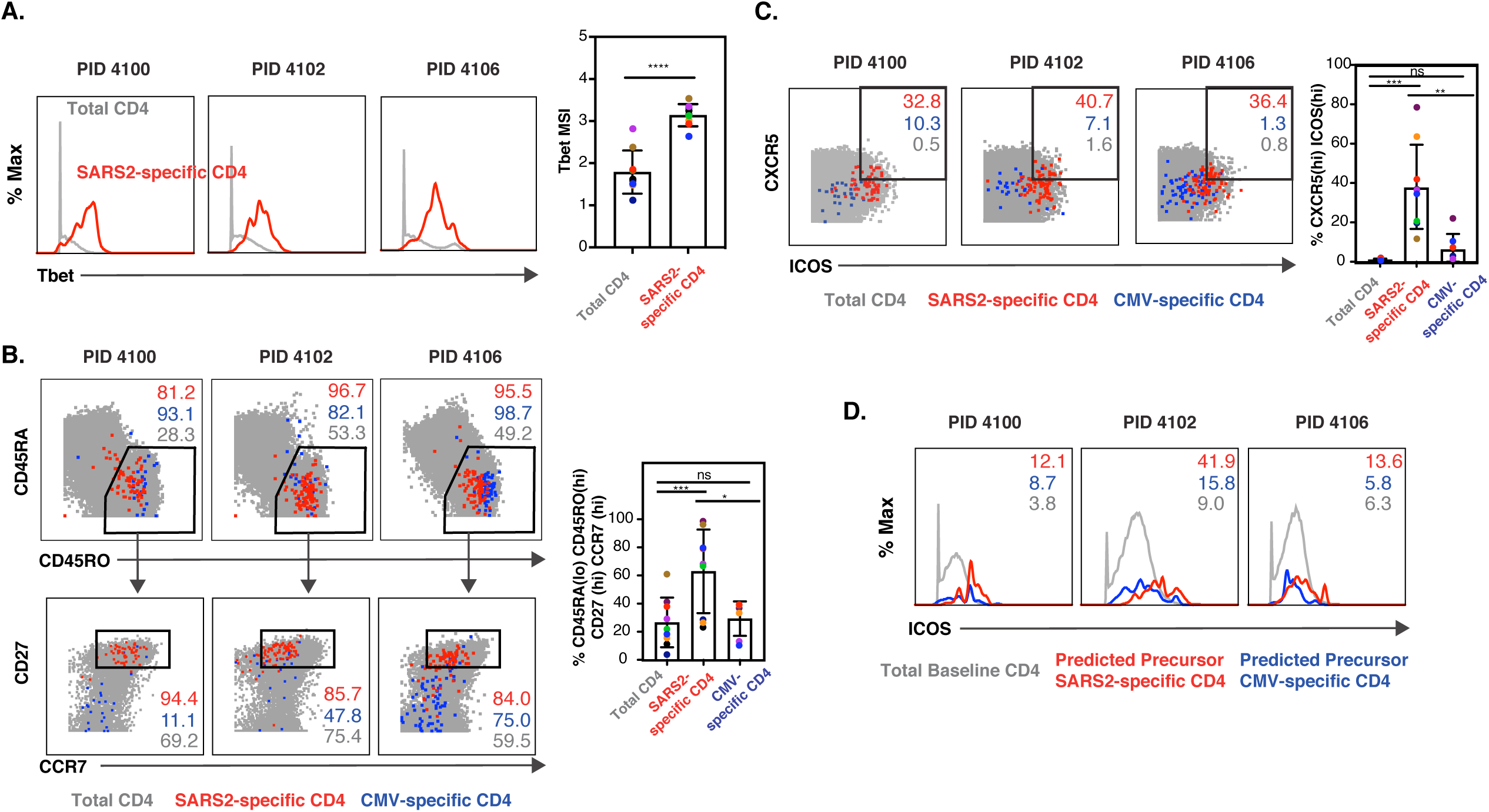
SARS-CoV-2-specific CD4+ Th1 cells are Tcm and cTfh. **A)** SARS-CoV2-specific CD4+ T cells are Th1 cells. The expression levels of Tbet, a transcription factor that directs Th1 differentiation, in total (*grey*) or SARS-CoV2-specific (*red*) CD4+ T cells from the blood of 3 representative convalescent individuals. Shown on the right are cumulative data from all 9 convalescent individuals analyzed in this study. **** p < 0.0001 as assessed using the Student’s paired t test. **B)** SARS-CoV-2-specific but not CMV-specific CD4+ T cells are predominantly Tcm. The phenotypes of total (*grey*), SARS-CoV-2-specific (*red*), and CMV-specific (*blue*) CD4+ T cells are shown as dot plots for 3 representative donors. *Top*: Both SARS-CoV-2-specific and CMV-specific CD4+ T cells are predominantly CD45RA-CD45RO+, characteristic of canonical memory cells. *Bottom:* Most memory (CD45RA-CD45RO+) SARS-CoV-2-specific CD4+ T cells are CD27+CCR7+, characteristic of Tcm, while most CMV-specific memory CD4+ T cells are CD27-CCR7-, characteristic of Tem. The percent of total, SARS-CoV-2-specific, and CMV-specific cells within the indicates gates are shown in grey, red, and blue, respectively. Shown on the right are cumulative data from all 9 convalescent individuals analyzed in this study. * p < 0.05, *** p < 0.001 as assessed using the Student’s unpaired t test. **C)** SARS-CoV-2-specific CD4+ T cells express high levels of CXCR5 and ICOS relative to total and CMV-specific CD4+ T cells. Numbers correspond to the percentages of SARS-CoV-2-specific (*red*), CMV-specific (*blue*), and total (*grey*) CD4+ T cells in the gates for 3 representative donors. Shown on the right are cumulative data from all 9 convalescent individuals analyzed in this study. ** p < 0.01, *** p < 0.001 as assessed using the Student’s unpaired t test. **D)** ICOS is expressed at high levels on predicted precursors of IFNγ-producing SARS-CoV-2-specific CD4+ T cells. PP-SLIDE (Cavrois et al., 2017; Ma et al., 2020b) was conducted to predict the original phenotypic features of SARS-CoV-2-specific (*red*) and CMV-specific (*blue*) cells prior to IFNγ induction. Expression levels of ICOS on these cells were compared to those on total CD4+ T cells phenotyped by CyTOF immediately following PBMC isolation. Numbers correspond to mean signal intensity (MSI) of ICOS expression for the populations indicated on the bottom.

ICOS serves a critical role in Tfh function (Hutloff et al., 1999), but is also an activation marker; therefore one concern was its levels were high on SARS-CoV-2-specific cells simply because these cells were responding to antigen stimulation. We believe that to not be the case because 1) ICOS-CXCR5-cells do not upregulate either ICOS or CXCR5 during 6 hours of *in vitro* stimulation (Bentebibel et al., 2013; Morita et al., 2011), and 2) SARS-CoV-2-specific CD4+ T cells had higher ICOS expression than CMV-specific CD4+ T cells which were similarly stimulated. To verify that SARS-CoV-2-specific cells at baseline express high levels of ICOS, we implemented Predicted Precursor as determined by SLIDE (PP-SLIDE) (Cavrois et al., 2017; Ma et al., 2020b), a bioinformatics pseudotime analysis approach that can predict the original phenotypes of cells before cellular perturbations. SARS-CoV-2- and CMV-specific CD4+ T cells were traced back to their predicted original states by matching their high-dimensional CyTOF profiles against the “atlas” of all CD4+ T cells phenotyped by CyTOF at baseline (prior to the 6 hours of stimulation). The predicted original states of SARS-CoV-2 had high levels of ICOS, supporting the notion that these cells exhibit phenotypic features of cells with robust helper function (Fig. 3D).

We next assessed whether SARS-CoV-2-specific CD4+ T cells exhibit features denoting longevity and an ability to proliferate. CD127, the alpha chain of the IL7 receptor, is involved in cell survival and is required for IL7-driven homeostatic proliferation (Kondrack et al., 2003). We found that among the nine convalescent donors, on average 58.5% +/- 20.5% of SARS-CoV-2-specific CD4+ T cells expressed CD127. Although the vast majority of CMV-specific CD4+ T cells also expressed CD127, these cells differed from their SARS-CoV-2-specific counterparts in that a higher proportion additionally expressed high levels of the terminal differentiation marker CD57 (Fig. 4A). To assess whether CD127+ SARS-CoV-2-specific CD4+ T cells are maintained over time, we conducted a phenotypic analysis of these cells in longitudinal specimens from two participants. SARS-CoV-2-specific CD4+ T exhibited stable phenotypes over time, and were detected more than two months post-infection (Fig. 4B). The proportions of CD127+ SARS-CoV-2-specific CD4+ T cells did not decrease over time, and in fact tended to increase (Fig. 4C).

**Figure 4.**
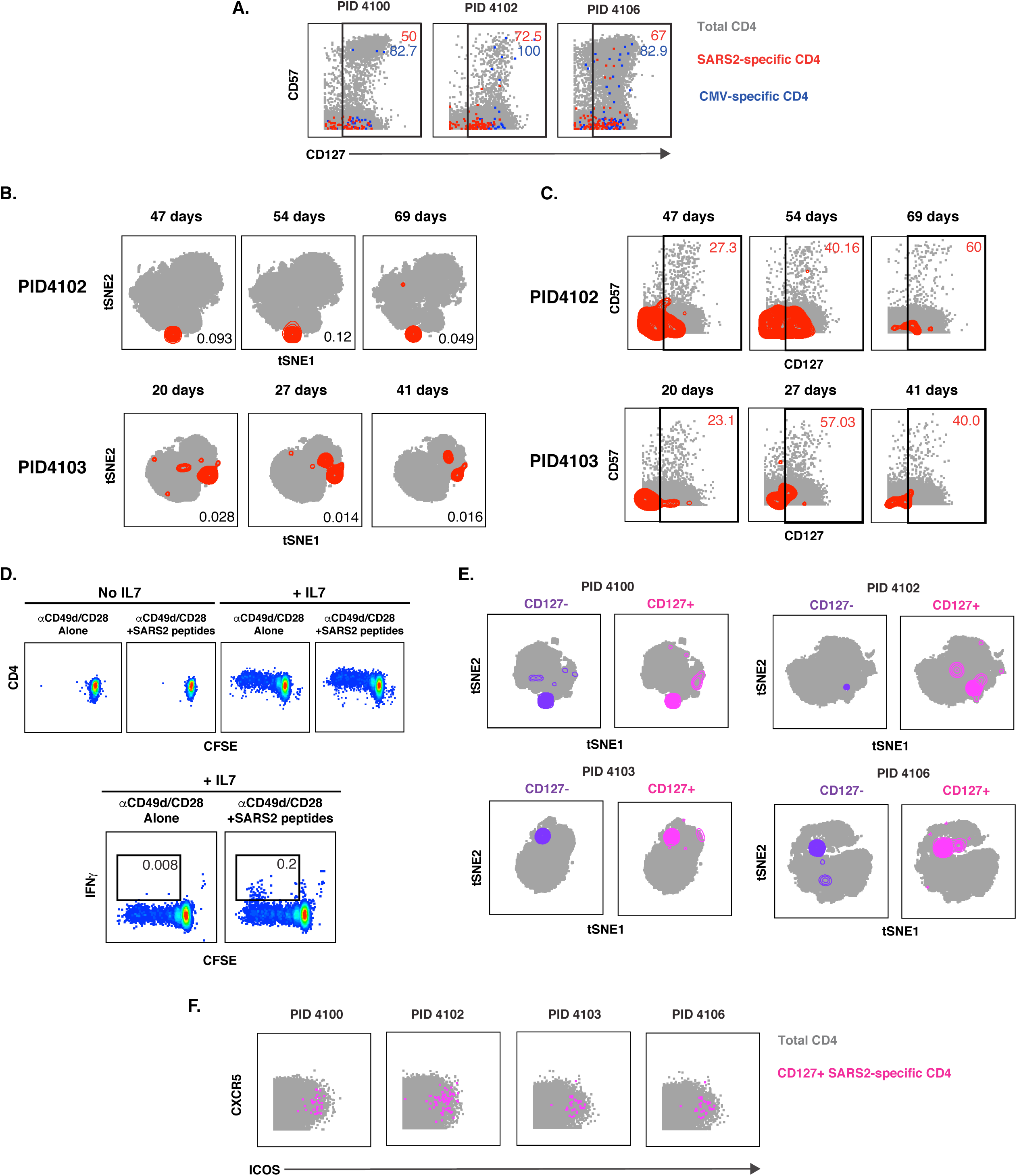
SARS-CoV-2-specific CD4+ T cells express CD127 and can persist for over 2 months. **A)** A subset of SARS-CoV-2-specific CD4+ T cells express CD127. The indicated cell populations were examined for expression levels of the terminal differentiation marker CD57 and the IL7 receptor CD127. Numbers correspond to the percentages of the corresponding populations within the gates. **B)** SARS-CoV-2-specific CD4+ T cells can persist for over two months. Shown are t-SNE plots of CyTOF datasets of CD4+ T cells from two COVID-19 convalescent donors that were sampled longitudinally at the indicated timepoints post-infection (infection defined as time of testing positive for SARS-CoV-2). Cells shown in grey correspond to total CD4+ T cells from specimens stimulated with anti-CD49d/CD28 in the absence of any peptides. The location of the SARS-CoV-2-specific cells are shown as red contours. The percent of CD4+ T cells that are SARS-CoV-2-specific is indicated in the bottom right of each plot. **C)** Persistent SARS-CoV-2-specific CD4+ T cells retain CD127 expression. Longitudinal specimens characterized in *panel B* were analyzed for expression levels of CD57 and CD127. SARS-CoV-2-specific CD4+ T cells are shown as red contours, while total CD4+ T cells are shown as grey dots. The percent of SARS-CoV-2-specific CD4+ T cells in the gates is shown in the top right of each plot. Note that the proportions of CD127+ SARS-CoV-2-specific CD4+ T cells do not decrease over time. **D)** SARS-CoV-2-specific CD4+ T cells can homeostatically proliferate in response to IL7. PBMCs from convalescent donor PID4102 were labeled with CFSE and then cultured for 5 days in the absence or presence of IL7, after which the cells were treated for 6 hours with co-stimulation alone or in the presence of overlapping peptides from SARS-CoV-2 spike, and then analyzed by flow cytometry. Homeostatic proliferation, as assessed by CFSE dye dilution, only occurred in the presence of IL7 (*top*). A sub-population of CFSE^low^ cells produced IFNγ in response to peptide stimulation (*bottom*), demonstrating that cells driven to proliferate by IL7 treatment included SARS-CoV-2-specific CD4+ T cells. Results are gated on live, singlet, CD3+CD4+CD8-cells, and are representative of one of two donors. **E)** CD127+ SARS-CoV-2-specific CD4+ T cells are similarly distributed as their CD127-counterparts. Shown are t-SNE plots of CD127-(*purple*) and CD127+ (*pink*) SARS-CoV-2-specific CD4+ T cells overlaid on total CD4+ T cells treated with co-stimulation alone (*grey*). Note that for each donor, the CD127+ cells occupy similar regions of t-SNE space as the CD127-cells. **F)** CD127+ SARS-CoV-2-specific CD4+ T cells express CXCR5 and ICOS.

To directly assess whether SARS-CoV-2-specific T cells were capable of homeostatic proliferation, we conducted an *in vitro* proliferation assay. PBMCs from convalescent donors were labeled with the proliferation dye CFSE, and then cultured for 5 days in the absence or presence of IL7. Treatment with IL7 induced proliferation of the cells, as reflected by dilution of the CFSE dye in a subset of the cells (Fig. 4D, *top*). After the 5 days of culture, the cells were subjected to intracellular cytokine staining analysis, following treatment with co-stimulation alone, or in the presence of SARS-CoV-2 spike peptides. SARS-CoV-2-specific CD4+ T cells were readily detected among CFSE^low^ CD4+ T cells (Fig. 4D, *bottom*), demonstrating that the cells that had homeostatically proliferated in response to IL7 treatment included SARS-CoV-2-specific CD4+ T cells.

To assess the phenotypic features of the CD127+ SARS-CoV-2-specific CD4+ T cells, we compared them to their CD127-counterparts. These two populations of cells occupied similar regions of t-SNE space (Fig. 4E), suggesting that the CD127+ and CD127-cells are phenotypically similar. Of note, the CD127+ cells include cells expressing high levels of CXCR5 and ICOS (Fig. 4F). Aside from CD127, CXCR4 was the only antigen that was expressed at significantly higher levels (p = 0.02 by pairwise test) on the CD127+ versus CD127-SARS-CoV-2-specific CD4+ T cells; however this difference was no longer significant when adjusted for multiple comparison using the Benjamini-Hochberg procedure (data not shown), consistent with the notion that the CD127+ and CD127-SARS-CoV-2-specific CD4+ T cells exhibit similar phenotypes.

In summary, SARS-CoV-2-specific CD4+ T cells from convalescent individuals exhibit phenotypic features consistent with an ability to migrate into lymph node follicles, provide robust helper function for B cells, and to be long-lived. A global view of all antigens differentially expressed in SARS-CoV-2-specific CD4+ T cells is presented in Fig. S6.

### SARS-CoV-2-specific CD8+ T cells exhibit phenotypic features of less-differentiated CD8+ Temra cells and express CD127

We next characterized the phenotypic features of SARS-CoV-2-specific CD8+ T cells. These cells, in stark contrast to their CD4+ counterparts, included a prominent population of CD45RA-expressing cells (Fig. 5A). CD45RA+ T cells include naïve (Tn) cells, but also Temra and stem cell memory (Tscm) cells. To determine the nature of the CD45RA-expressing CD8+ T cells, we assessed the proportions of CD8+ T cells that belonged to the Tn/Tscm (CD45RA+CD45RO-CCR7+) and Temra (CD45RA+CD45RO-CCR7-) subsets, as well as the canonical memory subsets Tcm (CD45RA-CD45RO+CCR7+CD27+), T effector memory (Tem) (CD45RA-CD45RO+CCR7-CD27-), and transitional memory (Ttm) (CD45RA-CD45RO+CCR7-CD27+). We then compared the subset distribution of the SARS-CoV-2-specific cells to those specific for CMV. Consistent with prior studies, CMV-specific CD8+ T cells were predominantly Temra cells (Derhovanessian et al., 2011) (Fig. 5B). Temra cells were also the largest subset of the SARS-CoV-2-specific CD8+ T cells (Fig. 5B). As CD8+ T cells recognizing influenza (flu) were also detected in a couple of the convalescent individuals, we compared their features to those recognizing SARS-CoV-2. Flu-specific and SARS-CoV-2 CD8+ T cells occupied different regions of the t-SNE, suggesting they are phenotypically distinct (Fig. S7A). While in one donor the flu-specific CD8+ T cells were not predominantly Temra cells, in the second they were (Fig. S7B, C). The variability in the phenotypes of the flu-specific T cells may be due to different times of flu exposure in these convalescent individuals.

**Figure 5.**
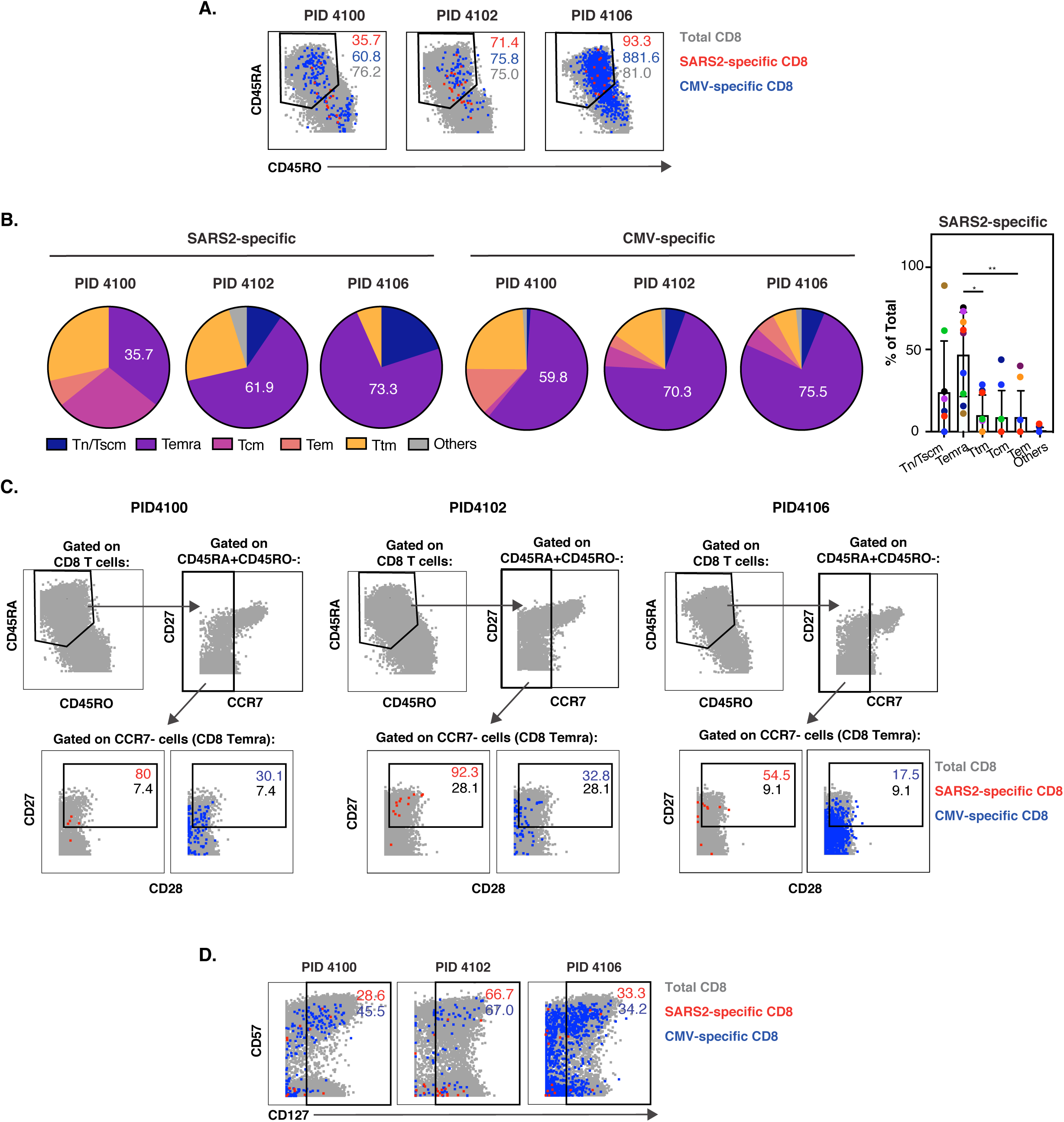
SARS-CoV-2-specific CD8+ T cells are predominantly a less differentiated subset of Temra and include long-lived CD127-expressing cells. **A)** SARS-CoV2-specific CD8+ T cells include both CD45RA+CD45RO- and CD45RA-CD45RO+ cells. The phenotypes of total (*grey*), SARS-CoV-2-specific (*red*), and CMV-specific (*blue*) CD8+ T cells are shown as dot plots for three representative donors. **B)** SARS-CoV-2-specific CD8+ T cells are predominantly Temra cells. The proportions of SARS-CoV-2-specific and CMV-specific CD8+ T cells belonging to each subset are depicted as pie graphs. Numbers correspond to the percentages of cells belonging to the Temra subset. Shown on the right are cumulative data for the SARS-CoV-2-specific CD8+ T cells from all 9 convalescent individuals analyzed in this study. * p < 0.05, * p < 0.01 as assessed using the Student’s paired t test. **C)** Compared to CMV-specific and total CD8+ Temra cells, SARS-CoV-2-specific CD8+ Temra cells express high levels of CD27 and CD28. *Top:* Temra cells were defined as CD45RA+CD45RO-cells expressing low levels of CCR7. *Bottom*: CD8+ Temra cells were further assessed for expression levels of CD27 and CD28. Numbers correspond to the percentage of total (*grey*), SARS-CoV-2-specific (*red*), and CMV-specific (*blue*) cells within the gate. **D)** A subset of SARS-CoV-2-specific CD8+ T cells express CD127. The indicated cell populations were examined for expression levels of CD57 and CD127.

To better define the features of the SARS-CoV-2-specific CD8+ Temra cells, and to compare them to their CMV-specific counterparts, we gated on these cells and assessed for expression levels of CD27 and CD28. These two co-stimulatory receptors have been used to distinguish the most terminally differentiated Temra cells (CD27-CD28-) from less differentiated ones (CD27+CD28+) (Koch et al., 2008). The vast majority of total CD8+ Temra cells in all donors expressed low levels of CD27, consistent with prior reports (Derhovanessian et al., 2011), while the majority of SARS-CoV-2-specific CD8+ Temra cells were CD27+ (Fig. 5C), suggesting that the SARS-CoV-2-specific CD8+ Temra cells were less terminally differentiated than typical Temra cells. These less differentiated CD8+ Temra cells were also present among CMV-specific cells, but in markedly lower proportions (Fig. 5C). Although CD28 staining was weak, it nonetheless suggested that CD28 levels were higher among the SARS-CoV-2-specific CD8+ T cells than among total or CMV-specific CD8+ T cells. These results suggest that although a large proportion of SARS-CoV-2-specific CD8+ T cells are Temra cells, they exhibit features characteristic of a less terminally differentiated state possibly capable of expansion.

Finally, to determine whether SARS-CoV-2-specific CD8+ T cells, like their CD4+ counterparts, may be capable of homeostatic proliferation, we assessed for expression levels of CD127. Although a lower proportion of the SARS-CoV-2-specific CD8+ T cells expressed CD127 relative to their CD4+ counterparts, when averaged over the nine convalescent donors, these cells still accounted for an average of 58.0% +/- 25.5% of the population. These CD127+ cells were similar to their CMV counterparts in that they included both CD57- and CD57+ cells (Fig. 5D). These results suggest that a subset of SARS-CoV-2-specific CD8+ T cells express CD127 and may therefore be long-lived. A global view of all antigens differentially expressed in SARS-CoV-2-specific CD8+ T cells is presented in Fig. S6.

## DISCUSSION

In this study, we define phenotypic features the T cell response against SARS-CoV-2 in nine convalescent individuals that recovered from COVID-19 after only experiencing mild symptoms. SARS-CoV-2-specific responses were readily detected in both CD4+ and CD8+ T cells from all nine individuals. The phenotypes of the SARS-CoV-2-specific CD4+ T cells were markedly different from those of their CD8+ counterparts, and included long-lived cells capable of homeostatic proliferation.

Recent studies measuring the levels of cytokines in supernatants of convalescent COVID-19 PBMCs treated with SARS-CoV-2 peptides revealed upregulation of IFNγ, suggesting a Th1 CD4+ T cell response (Grifoni et al., 2020; Weiskopf et al., 2020). However, in those studies the cellular source of the IFNγ was not determined. We confirmed the existence of SARS-CoV-2-specific CD4+ Th1 cells by demonstrating that Tbet-expressing CD4+ T cells from convalescent individuals secrete IFNγ in response to SARS-CoV-2 peptide treatment. These results suggest that similar to what has been observed for SARS-CoV (Janice Oh et al., 2012), a Th1 response directed against SARS-CoV-2 is associated with effective resolution of infection and symptoms. In comparison, the lack of any detectable SARS-CoV-2-specific CD4+ T cells producing IL4 or IL17 suggests that antigen-specific Th2 or Th17 responses are not key for recovery from COVID-19. In fact, Th2 and Th17 responses have been suggested to be detrimental for recovery from highly pathogenic coronavirus infections – Th2 responses due to their association with lung immunopathology in the context of SARS-CoV (Deming et al., 2006; Yasui et al., 2008), and Th17 responses due to the close associations of ARDS (exacerbated by Th17 responses) and IL6 (a Th17-associated cytokine) with severe COVID-19 (Hotez et al., 2020; Pacha et al., 2020). Interestingly, while we did not detect any IL6 production by SARS-CoV-2-specific CD4+ T cells (whereas IL6 induction was readily detectable upon LPS stimulation of monocytes, data not shown), IL6+ CD4+ T cells were reported to be elevated during severe COVID-19 (Zhou et al., 2020). Future studies are warranted to determine if SARS-CoV-2 CD4+ T cells from severely ill patients, particularly those in the ICU or those who succumb to disease, produce IL4, IL17, or IL6.

One important function of CD4+ T cells is to help B cells undergo affinity maturation, a process that typically occurs in lymph nodes draining the site of infection. We found that most SARS-CoV-2-specific CD4+ T cells were Tcm, cells poised for entering lymph nodes by virtue of their expressing the lymph-node homing chemokine receptor CCR7. Furthermore, we found that most SARS-CoV-2-specific CD4+ T cells expressed CXCR5, a chemokine receptor that directs Tfh into the germinal centers of lymph nodes where the B cells mature (Breitfeld et al., 2000; Schaerli et al., 2000). CXCR5 has been used as a marker of cTfh, the blood counterpart of lymphoid Tfh (Chevalier et al., 2011). Some cTfh also express high levels of ICOS, a co-stimulatory molecule that plays a critical role in Tfh function including helping the development of high-affinity B cells (Hutloff et al., 1999). We found that SARS-CoV-2-specific CD4+ T cells expressed especially high levels of ICOS, suggesting active helper function of B cells. These findings suggest that SARS-CoV-2-specific cTfh play a key role in effective immunity, and are in line with the observation that the frequency of total cTfh cells increases at the time of SARS-CoV-2 clearance (Thevarajan et al., 2020) and the detection of robust Tfh responses following exposure of rhesus macaques to SARS-CoV-2 (Elzaldi et al., 2020). Interestingly, the SARS-CoV-2-specific cTfh cells we identified exhibit features similar to a population of cTfh shown to be important in the generation of vaccine-induced antibodies against influenza (Bentebibel et al., 2013). Both express high levels of CXCR5 and ICOS. Both also appear to be Th1 cells – ours as defined by expression of Tbet and IFNγ, and the flu-responsive ones as defined by expression of CXCR3, a chemokine receptor preferentially expressed on Th1 cells (Bonecchi et al., 1998; Sallusto et al., 1998). Interestingly, the flu-responsive cTfh cells were shown to effectively help memory but not naïve B cells (Bentebibel et al., 2013). This, together with the observation that the frequencies of the Th1 cTfh cells were associated with flu-specific antibody titers only in individuals with pre-existing antibodies against other flu strains (Bentebibel et al., 2013), suggest these cells may be most effective at helping pre-existing cross-reactive memory B cells. Whether this is also true in the context of COVID-19 is not clear, but given that 40-60% of unexposed individuals have T cells that react against SARS-CoV-2 peptides, likely due to cross-reactivity with endemic coronaviruses (Grifoni et al., 2020), it is possible that the SARS-CoV-2-specific CD4+ T cells we identified are acting on previously-generated cross-reactive memory B cells. We speculate that the convalescent individuals we analyzed had previously been exposed to endemic coronaviruses and that this prior exposure, along with conditions favoring a Th1 response in these individuals, provided the necessary conditions for eliciting a robust and effective response against SARS-CoV-2. Testing this intriguing hypothesis will require serological and immune cell analyses of a large collection of specimens collected before and after exposure to SARS-CoV-2.

In contrast to their CD4+ counterparts, SARS-CoV-2-specific CD8+ T cells were predominantly Temra cells, antigen-experienced cells that re-express the naïve cell marker CD45RA. Due to lack of CCR7 expression, CD8+ Temra cells do not home efficiently to lymph nodes, and instead predominantly reside in blood, spleen, and lung (Thome et al., 2014). These cells have been shown to be protective against a variety of viral pathogens, including CMV, flu, EBV, and HIV (Dunne et al., 2002; Lilleri et al., 2008; Northfield et al., 2007; Sridhar et al., 2013). Exceptionally high numbers of CMV-specific Temra have been observed in CMV-infected individuals (van den Berg et al., 2019), and this subset also constitutes a major proportion of dengue-specific CD8+ T cells in individuals that have recovered from dengue infection (Chng et al., 2019; Tian et al., 2019). Although generally thought to be terminally differentiated, some CD8+ Temra can be long-lived. The most terminally differentiated CD8+ Temra cells express low levels of CD27 and CD28, and these cells are thought to have differentiated from a more pluripotent state of Temra cells expressing both CD27 and CD28 (Koch et al., 2008).

Interestingly, SARS-CoV-2-specific CD8+ Terma were heavily biased towards the CD27+CD28+ subset, when compared to both total as well as CMV-specific CD8+ Temra. These phenotypic features are similar to the previously-described EBV-specific CD8+ Temra cells, a subset that was demonstrated to be apoptosis-resistant due to expression of bcl2, to have long telomere lengths relative to other memory subsets (suggesting high replicative potential), and to be capable cytotoxic function (Dunne et al., 2002). NK-like cytotoxic functions have also been observed in dengue-specific CD8+ Temra (Tian et al., 2019). We postulate that based on their phenotypic similarities to EBV-specific CD8+ Temra cells, SARS-CoV-2-specific CD8+ Temra cells are cytotoxic and long-lived, although this will need to be confirmed in follow-up studies.

Consistent with the notion of long-lived cells, we found that a substantial fraction of both CD4+ and CD8+ T cells specific for SARS-CoV-2 expressed CD127. This surface protein corresponds to the alpha chain of the IL7 receptor, and together with the common gamma chain (CD132) shared by the IL-2 and IL-15 receptors, enables T cell proliferation in response to IL-7. Signaling by IL-7 is involved in many key aspects of T cell survival and proliferation, and acts through inducing JAK/STAT signaling which results in increased expression of the Bcl2 anti-apoptotic gene family (Kaech et al., 2003; Kondrack et al., 2003). Consistent with the notion of persisting CD127-expressing SARS-CoV-2-specific T cells is our observation that these cells could still be detected 69 days post-infection, and that they can proliferate in response to IL7. Collectively, these results suggest that SARS-CoV-2-specific T cell responses have the potential to be long-lived, something that has been found to be true for SARS-CoV where memory T cells persist for up to 11 years post-infection (Ng et al., 2016; Tang et al., 2011). Having long-lived protective T cells against SARS-CoV-2 may be particularly important as no neutralizing antibodies were detected in 18% of convalescent individuals ∼30 days after symptom onset, suggesting that the antibody response may be short-lived, at least in some individuals (Robbiani et al., 2020). Of note, a clinical trial (NCT04379076) has been initiated to treat hospitalized lymphopenic COVID-19 patient with IL7, in an effort to restore T cell counts (Gill and Buchanan, 2020). Whether such treatment will boost the numbers of SARS-CoV-2-specific T cells remains to be determined, but is conceivable given that CD127 is expressed on these cells.

In summary, our phenotypic analysis reveals long-lived lymph-node homing cTfh CD4+ T cells and CD27+CD28+ CD8+ Temra as the major subsets of SARS-CoV-2-specific T cells that persist following recovery from mild COVID-19. The system we developed here to conduct high-parameter phenotypic analysis of SARS-CoV-2-specific T cells can be used in future studies to better understand what constitutes an effective versus an immunopathological T cell response against the virus, and to assess the features of vaccine-induced SARS-CoV-2-specific T cell responses. As our study was limited to analyzing T cells from nine individuals who recovered efficiently from COVID-19, it does not allow us to determine exactly why some individuals fight off the infection while others succumb to it. Nonetheless, it revealed common features of effective immunity against SARS-CoV-2, and suggests that mobilization of a Th1 response plays a key role in viral clearance and the establishment of long-lived CD4+ and CD8+ T cells. Vaccination strategies aimed at conferring long-term COVID-19 immunity should strongly consider approaches that include elicitation of long-lived CD4+ and CD8+ T cells against this pandemic virus.

## AUTHOR CONTRIBUTIONS

J.N., J.F., G.X., and A.G. conducted experiments; J.N., X.L., M.M., T.M., A.K., E.G., and N.R.R.. conducted data analysis; X.L. and M.M. wrote scripts; J.N., J.F., G.X., E.G., and N.R.R. designed the experiments; S.L. established the CHIRP cohort; G.G., E.S.S., and S.L. conducted CHIRP participant interviews, enrollment, and specimen collection; W.C.G. supervised data analysis and edited the manuscript; J.V., E.G., S.L., and N.R.R. conceived ideas for the study; N.R.R. wrote the manuscript. All authors read and approved the manuscript.

## ACKNOWLEDGEMENTS

This work was supported by the Van Auken Private Foundation and David Henke; the Program for Breakthrough Biomedical Research, which is partly funded by the Sandler Foundation; and philanthropic funds donated to the Gladstone Institutes by The Roddenberry Foundation and individual donors devoted to COVID-19 research. We acknowledge the NIH DRC Center Grant P30 DK063720 and the S10 1S10OD018040-01 for use of the CyTOF instrument. We also thank Stanley Tamaki, Tomoko Kakegawa Peech, and Caudia Bispo for CyTOF assistance at the Parnassus Flow Core, Nicole Lazarus and Eugene Butcher for the Act1 antibody, Françoise Chanut for editorial assistance, and Robin Givens for administrative assistance.

## SUPPLEMENTARY FIGURE LEGENDS

**Figure S1.**
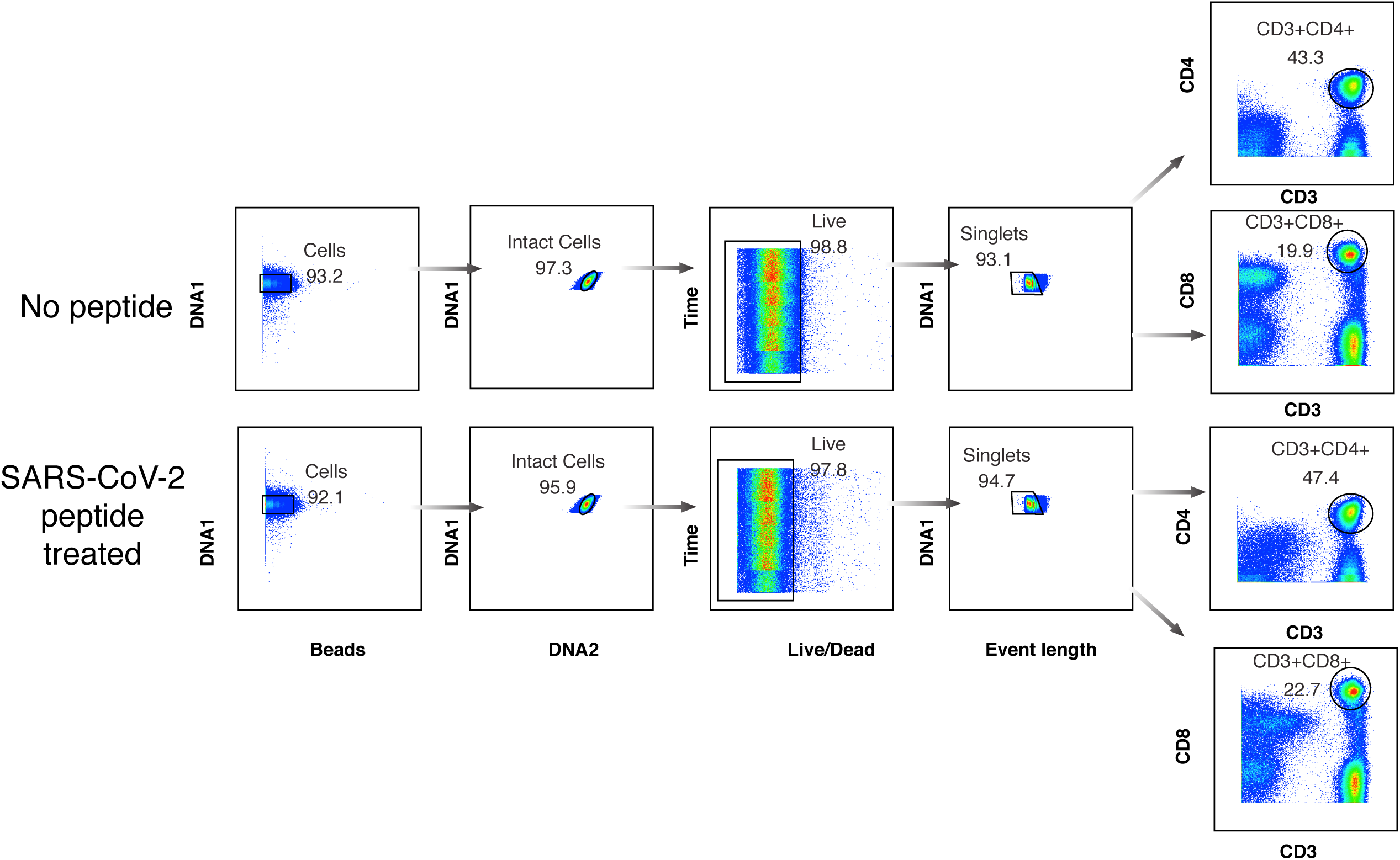
CyTOF gating strategy to identify CD4+ and CD8+ T cells from convalescent COVID-19 patients. PBMCs were purified from freshly drawn blood specimens, treated as indicated in Fig. 1, and phenotyped by CyTOF. Shown is an example of the gating strategy leading to the identification of live, singlet CD4+ and CD8+ T cells.

**Figure S2.**
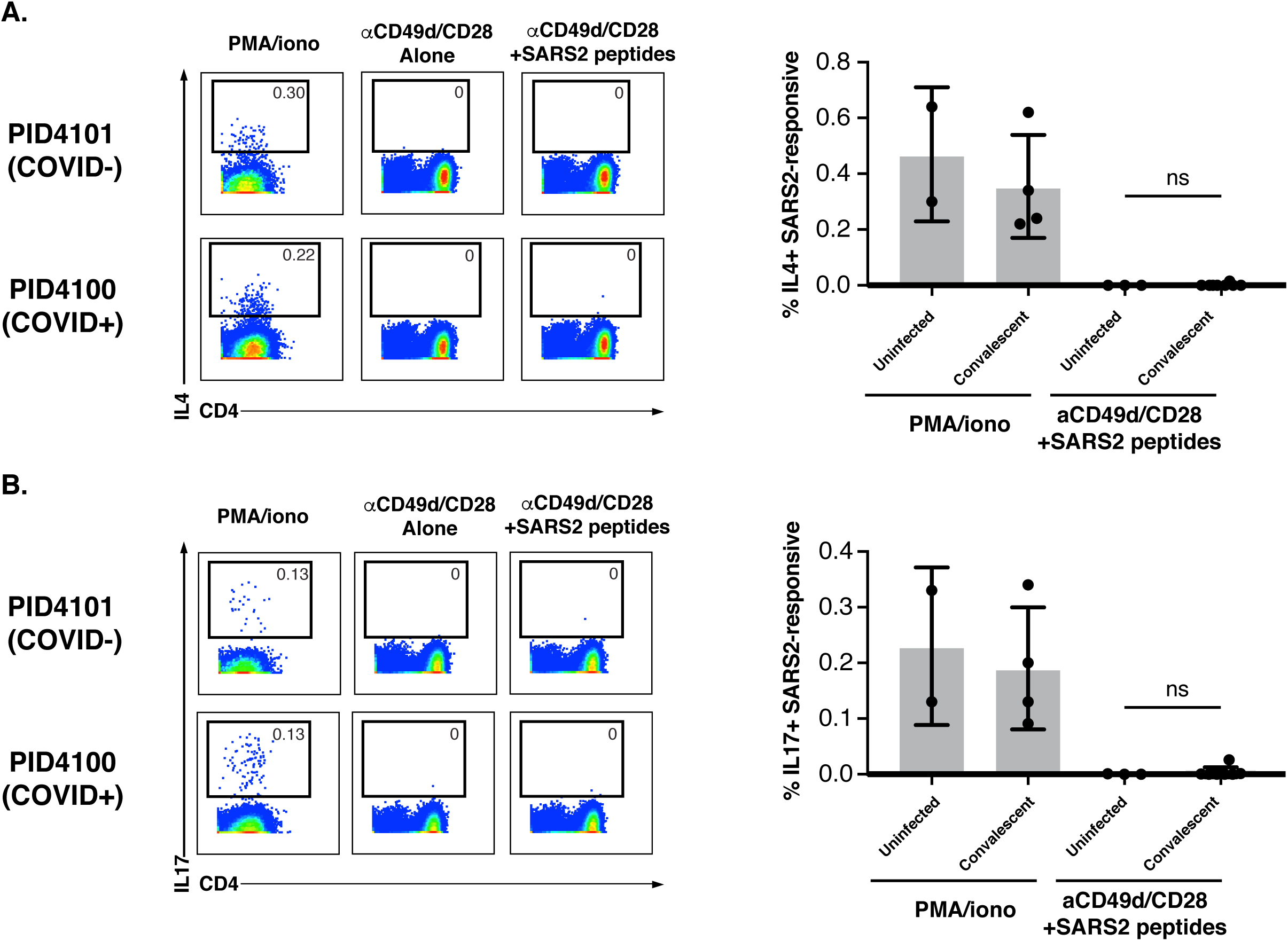
Blood from convalescent individuals lack SARS-CoV-2 spike-specific CD4+ T cells producing IL4 or IL17. Shown on the left are pseudocolor plots of CyTOF datasets reflecting the percentage of CD4+ T cells producing IL4 (**A**) or IL17 (**B**) in response to the indicated treatment condition, for one representative uninfected (COVID-) and one convalescent (COVID+) donor. Numbers correspond to the percentage of cells within the gates. PMA/ionomycin stimulation was used to demonstrate the presence of IL4- and IL17-producing T cells in these donors. Anti-CD49d/CD28 was used to provide co-stimulation during peptide treatment. Shown on the right are cumulative data of three uninfected individuals and nine convalescent individuals (Table S2). Results are gated on live, singlet CD4+ T cells. n.s. = non-significant as assessed using the Student’s unpaired t test.

**Figure S3.**
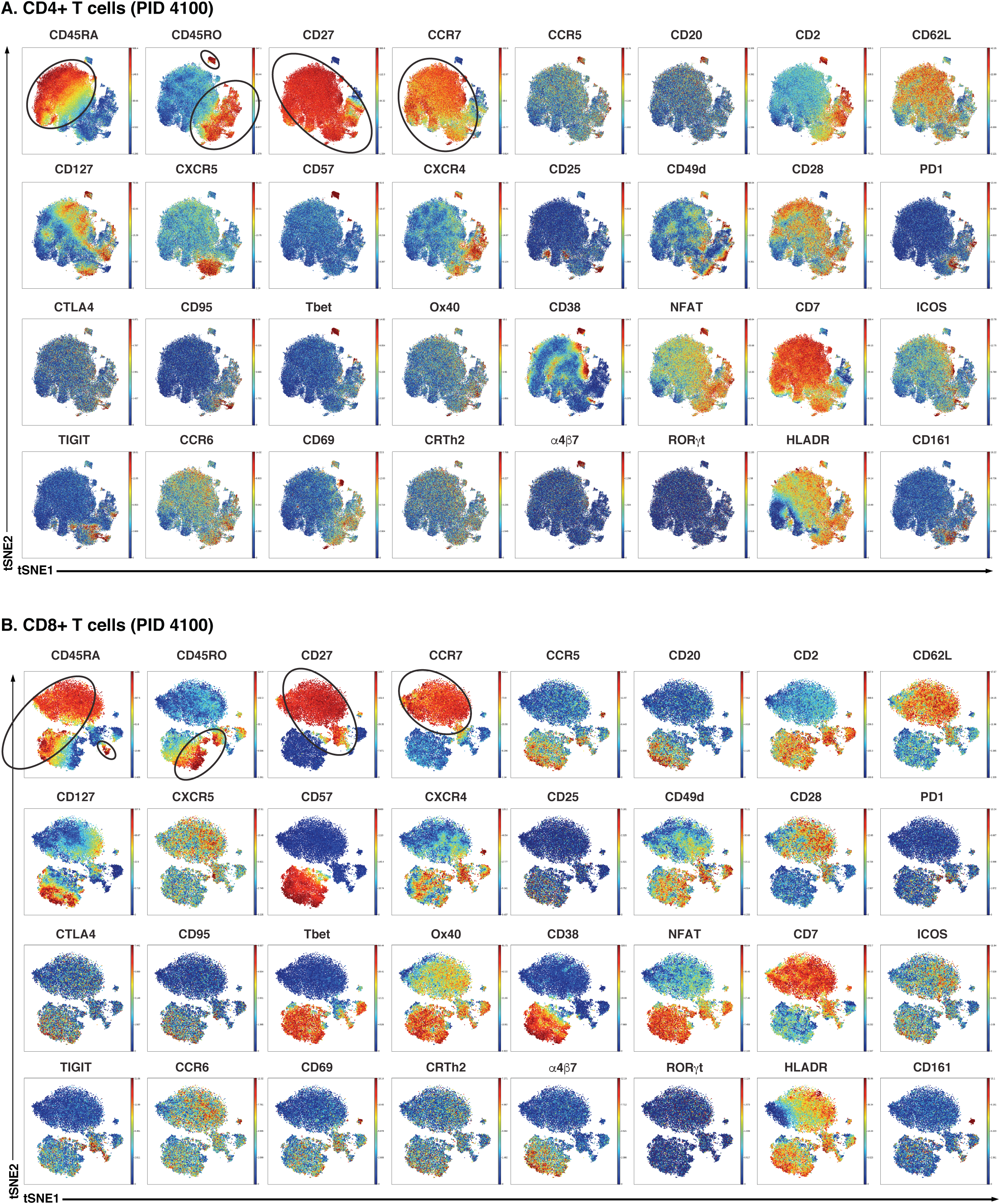
Heatmaps of antigen expression in T cells from representative donor. Shown are t-SNE plots of CyTOF datasets reflecting CD4+ (A) and CD8+ (B) T cells from representative convalescent donor PID4100. Regions of the t-SNE harboring SARS-CoV-2- and CMV-specific T cells are shown in Fig. 2. Circled in the first four plots are regions with high expression levels of CD45RA (expressed on naïve/Tscm/Temra T cells), CD45RO (expressed on memory T cells), and CD27 and CCR7 (expressed on naïve and Tcm cells).

**Figure S4.**
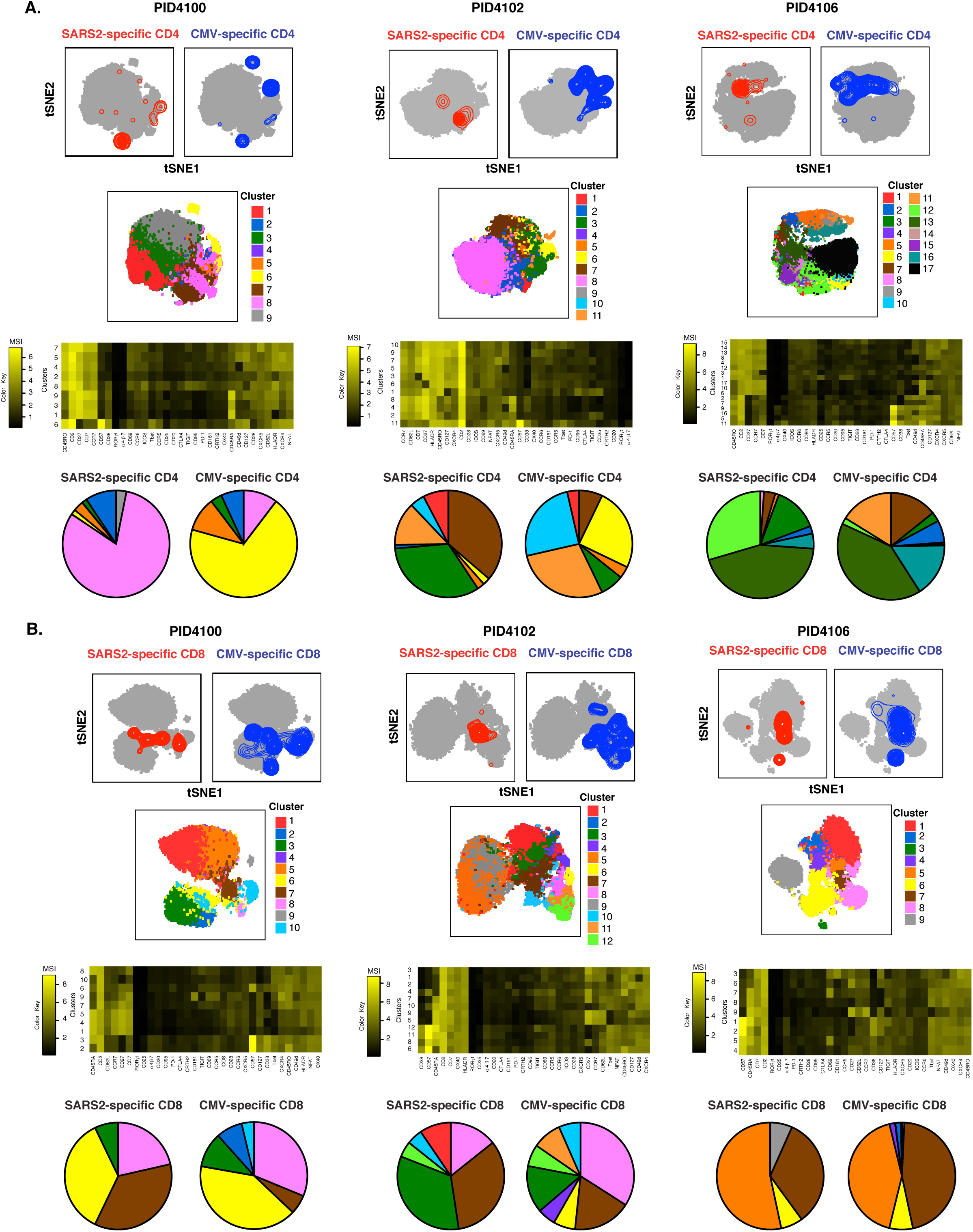
SARS-CoV-2 spike-specific T cells recognizing SARS-CoV-2 differ in phenotypes from those recognizing CMV. Shown are t-SNE plots of CyTOF datasets reflecting CD4+ (**A**) or CD8+ (**B**) T cells from three CMV+ COVID-19 convalescent donors. The top pairs of plots are identical to the contour plots presented in Fig. 2 and serve as references for where in the t-SNE the SARS-CoV-2-specific and CMV-specific T cells are concentrated. The middle plots depict the same t-SNE colored by DensVM clusters. Shown underneath the clustered t-SNE plots are heatmaps showing relative expression levels (in mean signal intensity, or MSI) for each of the indicated antigens, hierarchically clustered based on Euclidean distances. The pie graphs at the bottom depict the proportions of SARS-CoV-2-specific or CMV-specific T cells that belong to each cluster. Note that each pair of pie graphs differs, with more pronounced differences observed among the CD4+ T cells.

**Figure S5.**
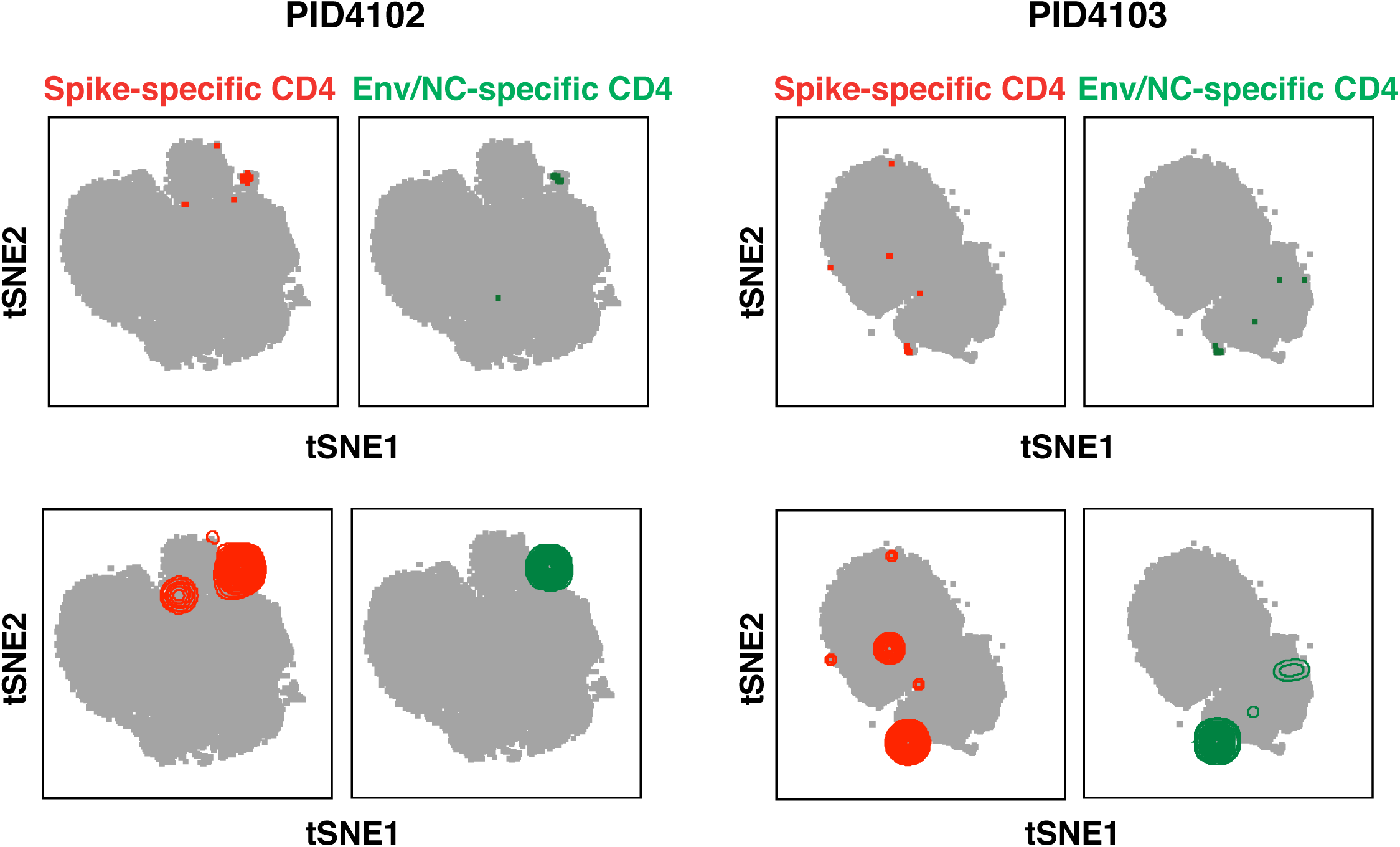
Antigen-specific CD4+ T cells against SARS-CoV-2 spike, envelope (env), and nucleocapsid (NC) are phenotypically similar. Shown are t-SNE plots of CyTOF datasets reflecting CD4+ T cells from two COVID-19 convalescent individuals. Cells shown in grey correspond to CD4+ T cells from specimens stimulated with anti-CD49d/CD28 in the absence of any peptides. The top pairs of plots show SARS-CoV-2 spike-specific (*red*) or env/NC-specific (*green*) cells as individual dots. The bottom pairs of plots show the same data but with the antigen-specific cells shown as contours instead of dots, to better visualize regions with highest densities of antigen-specific cells. Note that the highest densities of antigen-specific cells reside in similar regions of the t-SNE, suggesting that CD4+ T cells recognizing spike, env, and NC are phenotypically similar.

**Figure S6.**
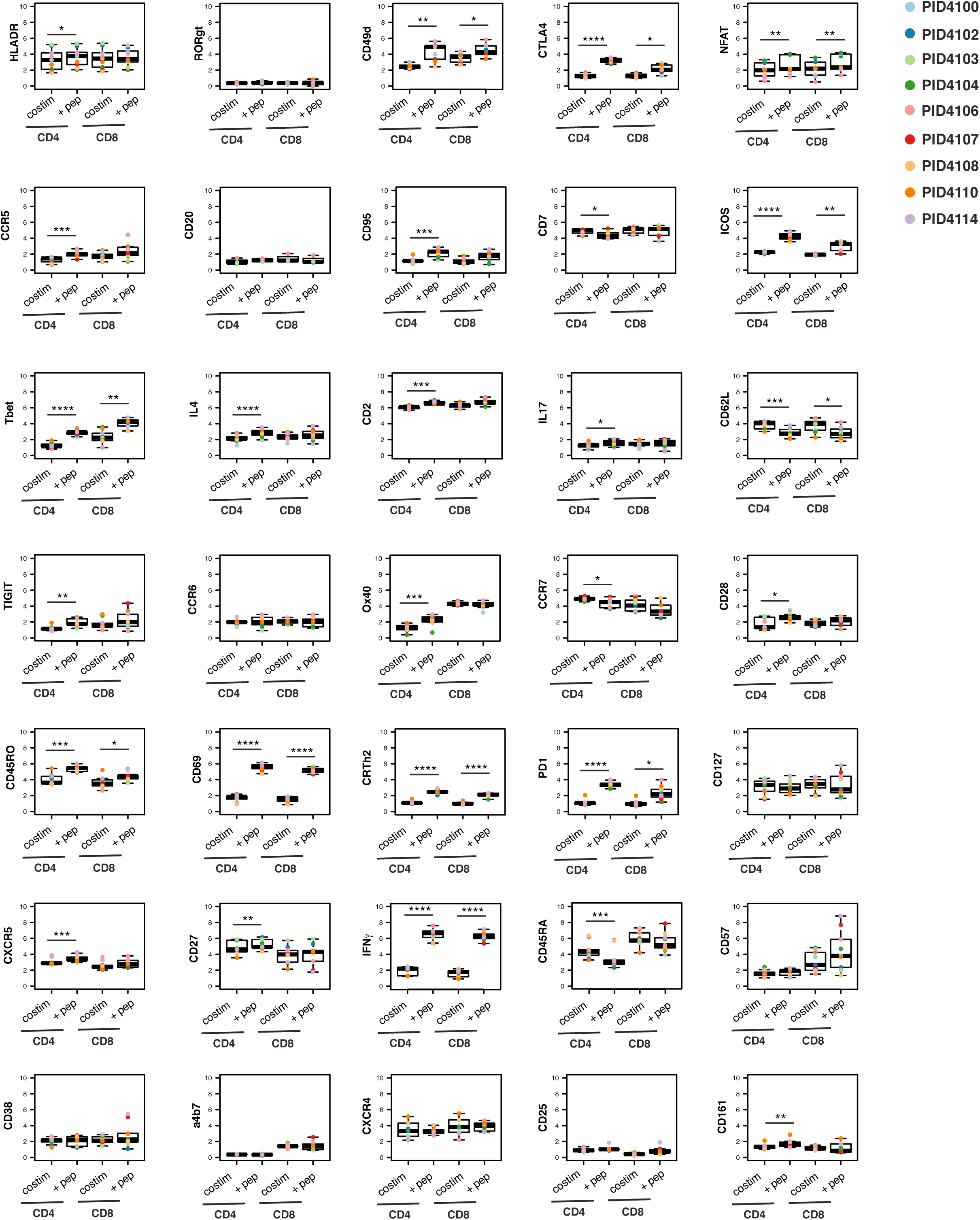
Mean expression levels of 35 surface and intracellular antigens in CD4+ and CD8+ T cells from 9 convalescent individuals who had recovered from mild COVID-19. PBMCs were purified from freshly drawn blood specimens, treated with anti-CD49d/CD28 for 6 hours alone (“costim”), or in the additional presence of overlapping 15-mer peptides against SARS-CoV-2 spike (“+ pep”), and then phenotyped by CyTOF. Results are gated on live, singlet CD4+/CD8+ T cells for the “costim” conditions, and live, singlet CD4+/CD8+ T cells expressing IFNγ for the “+ pep” conditions. Shown are the mean signal intensity (MSI) values for arcsinh-transformed data. *p < 0.05, **p < 0.01 as assessed using the Student’s paired t test and adjusted for multiple testing using the Benjamini-Hochberg for FDR.

**Figure S7.**
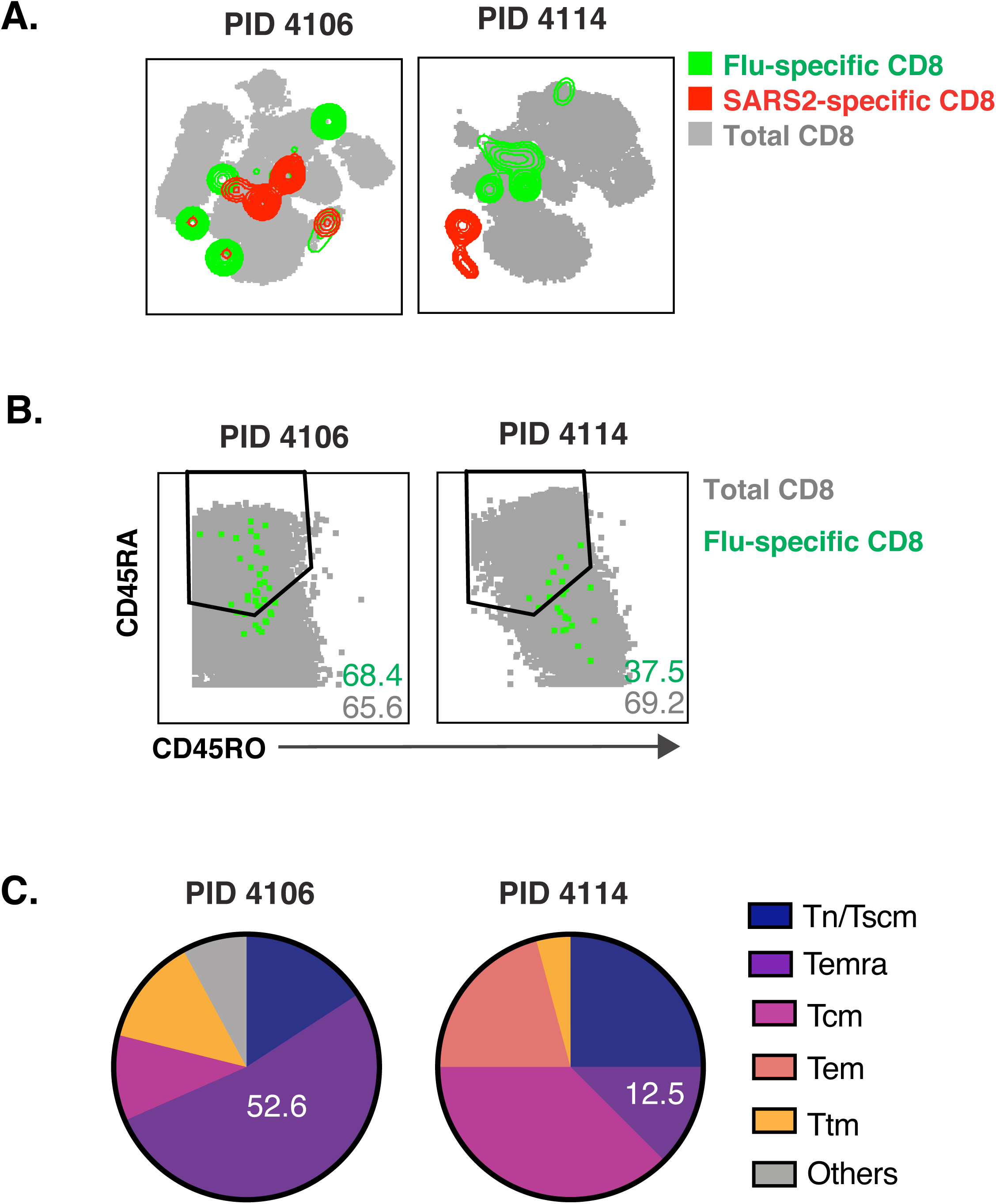
Flu-specific CD8+ T cells in convalescent individuals are phenotypically distinct from SARS-CoV-2-specific CD8+ T cells. **A)** Shown are t-SNE plots of CyTOF datasets reflecting CD8+ T cells from two COVID-19 convalescent donors who harbored flu-specific CD8+ T cell responses. Cells shown in grey correspond to CD8+ T cells from specimens stimulated with anti-CD49d/CD28 in the absence of any peptides. Flu- and SARS-CoV-2-specific CD8+ T cells shown as green and red contours, respectively. **B)** Flu-specific CD8+ T cells include both CD45RA+CD45RO- and CD45RA-CD45RO+ cells. The phenotypes of total (*grey*) or flu-specific (*green*) CD8+ T cells are shown as dot plots for two COVID-19 convalescent donors for which flu-specific responses were detected. **C)** Temra cells can comprise a majority or minority of the flu-specific CD8+ T cell response. The proportions of flu-specific CD8+ T cells belonging to each subset are depicted as pie graphs. Numbers correspond to the percentages of cells belonging to the Temra subset.

**Table S1:** Participant Characteristics

| <u>Patient ID</u> | <u>Biological Sex</u> | <u>Age</u> | <u>SARS-CoV-2 Status</u> | <u>CMV status</u> | <u>Date of symptom onset</u> | <u>Date of PCR+ test</u> | <u>Date of blood draw(s)</u> | <u>Time between PCR+ test and analysis</u> |
| --- | --- | --- | --- | --- | --- | --- | --- | --- |
| PID4100 | Female | 28 | Infected | Positive | 3/17/20 | 3/19/20 | 4/24/20 | 36 days |
| PID4102 | Male | 46 | Infected | Positive | 3/11/20 | 3/13/20 | 4/29/20<br>5/6/20*<br>5/20/20* | 47 days,<br>54 days*<br>69 days* |
| PID4103 | Female | 41 | Infected | Negative | 3/13/20 | 4/9/20 | 4/29/20<br>5/6/20*<br>5/20/20* | 20 days,<br>27 days*<br>41 days* |
| PID4104 | Female | 33 | Infected | Negative | 3/11/20 | 3/14/20 | 5/13/20 | 60 days |
| PID4106 | Female | 67 | Infected | Positive | 3/6/20 | 3/26/20 | 5/4/20 | 39 days |
| PID4114 | Female | 45 | Infected | Positive | 4/15/20 | 4/17/20 | 6/15/20 | 59 days |
| PID4107 | Female | 38 | Infected | Positive | 3/25/20 | 4/1/20 | 5/13/20 | 42 days |
| PID4108 | Female | 18 | Infected | Positive | 3/16/20 | NA | 5/13/20 | NA |
| PID4110 | Male | 57 | Infected | Positive | 4/19/20 | 4/24/20 | 5/15/20 | 21 days |
| PID4101 | Female | 42 | Uninfected | Positive | NA | NA | 4/24/20 | NA |
| PID4105 | Male | 57 | Uninfected | Negative | NA | NA | 5/4/20 | NA |
| PID4109 | Male | 30 | Uninfected | Negative | NA | NA | 5/13/20 | NA |

\*Used in longitudinal analysis

**Table S2:** List of CyTOF antibodies used in study. Antibodies were either purchased from the indicated vendor or prepared in-house using commercially available MaxPAR conjugation kits per manufacturer’s instructions (Fluidigm).

| **Antigen Target** | **Clone** | **Elemental Isotope** | **Vendor** |
| --- | --- | --- | --- |
| HLADR | TÜ36 | Qdot (112Cd) | Life Technologies |
| RORγt* | AFKJS-9 | 115 In | In-house |
| CD49d (α4) | 9F10 | 141Pr | Fluidigm |
| CTLA4* | 14D3 | 142Nd | In-house |
| NFAT* | D43B1 | 143Nd | Fluidigm |
| CCR5 | NP6G4 | 144Nd | Fluidigm |
| CD20 | 2H7 | 145Nd | In-house |
| CD95 | BX2 | 146Nd | In-house |
| CD7 | CD76B7 | 147Sm | Fluidigm |
| ICOS | C398.4A | 148Nd | Fluidigm |
| Tbet* | 4B10 | 149Sm | In-house |
| IL4* | MP4-25D2 | 150Nd | In-house |
| CD2 | TS1/8 | 151Eu | Fluidigm |
| IL17* | BL168 | 152Sm | In-house |
| CD62L | DREG56 | 153Eu | Fluidigm |
| TIGIT | MBSA43 | 154Sm | Fluidigm |
| CCR6 | 11A9 | 155Gd | In-house |
| IL6* | MQ2-13A5 | 156 Gd | In-house |
| CD8 | RPA-T8 | 157Gd | In-house |
| CD19 | HIB19 | 157Gd | In-house |
| CD14 | M5E2 | 157Gd | In-house |
| OX40 | ACT35 | 158Gd | Fluidigm |
| CCR7 | G043H7 | 159Tb | Fluidigm |
| CD28 | CD28.2 | 160Gd | Fluidigm |
| CD45RO | UCHL1 | 161Dy | In-house |
| CD69 | FN50 | 162Dy | Fluidigm |
| CRTH2 | BM16 | 163Dy | Fluidigm |
| PD-1 | EH12.1 | 164Dy | In-house |
| CD127 | A019D5 | 165Ho | Fluidigm |
| CXCR5 | RF8B2 | 166Er | In-house |
| CD27 | L128 | 167Er | Fluidigm |
| IFNg* | B27 | 168Er | Fluidigm |
| CD45RA | HI100 | 169Tm | Fluidigm |
| CD3 | UCHT1 | 170Er | Fluidigm |
| CD57 | HNK-1 | 171Yb | In-house |
| CD38 | HIT2 | 172Yb | Fluidigm |
| $\alpha 4\beta 7$ | Act1 | 173Yb | In-house |
| CD4 | SK3 | 174Yb | Fluidigm |
| CXCR4 | 12G5 | 175Lu | Fluidigm |
| CD25 | M-A251 | 176Yb | In-house |
| CD161 | NKR-P1A | 209 Bi | In-house |

\*Intracellular antibodies

**Table S3:** Frequencies of SARS-CoV-2- and CMV-specific T cells in convalescent individuals

| Patient ID | SARS2 CD4* | SARS2 CD8<sup>#</sup> | CMV CD4* | CMV CD8<sup>#</sup> |
| --- | --- | --- | --- | --- |
| PID4100 | 0.067 | 0.05 | 0.36 | 0.66 |
| PID4102 | 0.28 | 0.21 | 0.061 | 0.47 |
| PID4103 | 0.054 | 0.038 | 0 | 0 |
| PID4104 | 0.068 | 0.009 | 0 | 0 |
| PID4106 | 0.14 | 0.07 | 0.18 | 6.9 |
| PID4107 | 0.030 | 0.14 | 0.016 | 0.049 |
| PID4108 | 0.029 | 0.015 | 0.026 | 0.027 |
| PID4110 | 0.14 | 0.026 | 0.048 | 0.0052 |
| PID4114 | 0.076 | 0.010 | 0.13 | 0.037 |

\* Reported as frequency of live, singlet CD4+ T cells
<sup>#</sup> Reported as frequency of live, singlet CD8+ T cells

## METHODS

### Study Participants and Specimen Collection

Blood was obtained from eight individuals who had confirmed SARS-CoV-2 as assessed by RT-PCR, one household member of a confirmed SARS-CoV-2-positive individual who was not tested but exhibited COVID-19 symptoms (PID4107) and was therefore assumed to be infected, and three healthy controls. All participants were recruited as part of the UCSF acute COVID-19 Host Immune Pathogenesis (CHIRP) study. The timing of specimen collection relative to symptom onset and when SARS-CoV-2 infection was confirmed are presented in Table S1. Two individuals were additionally sampled at three longitudinal study visits. This study was approved by the University of California, San Francisco (IRB # 20-30588).

### Preparation of participant specimens for CyTOF

PBMCs were isolated from blood using Lymphoprep^™^ (StemCell Technologies) within 2 hours of blood collection. Six million cells were then immediately treated with cisplatin (Sigma-Aldrich) as a Live/Dead marker and fixed with paraformaldehyde (PFA) as previously described (Ma et al., 2020b). Briefly, 6×10^6^ cells were resuspended at room temperature in 2 ml PBS (Rockland) with 2 mM EDTA (Corning). Next, 2 ml of PBS containing 2 mM EDTA and 25 μM cisplatin (Sigma-Aldrich) were added to the cells. The cells were quickly mixed and incubated at room temperature for 60 seconds, after which 10 ml of CyFACS (metal contaminant-free PBS (Rockland) supplemented with 0.1% FBS and 0.1% sodium azide (Sigma-Aldrich)) was added to quench the reaction. The cells were then centrifuged and resuspended in 2% PFA in CyFACS, and incubated for 10 minutes at room temperature. The cells were then washed twice in CyFACS, after which they were resuspended in 100 μl of CyFACS containing 10% DMSO. These fixed cells were stored at -80°C until analysis by CyTOF.

For identification of antigen-specific T cells, unless otherwise indicated, 6 million freshly-isolated PBMCs were stimulated for 6 hours in RP10 media (RPMI 1640 medium (Corning) supplemented with 10% fetal bovine serum (FBS, VWR), 1% penicillin (Gibco), and 1% streptomycin (Gibco)) in the presence of 3 μg/ml Brefeldin A Solution (eBioscience) to enable detection of intracellular cytokines. For detection of antigen-specific T cells, 0.5 μg/ml anti-CD49d clone L25 and 0.5 μg/ml anti-CD28 clone L293 (both from BD Biosciences) were added as a source of co-stimulation, in the presence of 0.5 μm PepMix™ SARS-CoV-2 Peptide (Spike Glycoprotein) (JPT Peptide Technologies), human CMV pp65 Peptide Pool (NIH AIDS Reagent Program), or Influenza Virus Control Peptide Pool (Anaspec). As a positive control for cytokine detection, cells were treated with 16 nM PMA (Sigma-Aldrich) and 1 μm ionomycin (Sigma-Aldrich). Treated cells were then cisplatin-treated and PFA-fixed as described above. For longitudinal comparison of SARS-CoV-2-specific T cells, cryopreserved PBMCs from 3 timepoints each from participants PID4102 and PID4103 were simultaneously thawed, and then cultured overnight in RP10 media. Peptide stimulation was conducted for 6 hours as described above, using 0.5 μm PepMix™ SARS-CoV-2 Peptides for the following: spike glycoprotein, or a mix of envelope (env) and nucleocapsid (NC) (JPT Peptide Technologies), prior to cisplatin treatment and PFA fixation. Due to logistical reasons, PBMCs from participants PID4107, PID4108, and PID4109 were also cryopreserved prior to peptide stimulation and CyTOF analysis.

### CyTOF staining and data acquisition

Staining of cells for analysis by CyTOF was conducted similar to recently described methods (Cavrois et al., 2017; Ma et al., 2020a; Trapecar et al., 2017). Briefly, cisplatin-treated cells were thawed and washed in Nunc 96 DeepWell^™^ polystyrene plates (Thermo Fisher) with CyFACS buffer at a concentration of 6⨯10^6^ cells / 800 μl in each well. Cells were then pelleted and blocked with mouse (Thermo Fisher), rat (Thermo Fisher), and human AB (Sigma-Aldrich) sera for 15 minutes at 4°C. The samples were then washed twice in CyFACS, pelleted, and stained in a 100-μl cocktail of surface antibodies (Table S2) for 45 minutes at 4°C. The samples were then washed three times with CyFACS and fixed overnight at 4°C in 100 μl of freshly prepared 2% PFA in PBS (Rockland). Samples were then washed twice with Intracellular Fixation & Permeabilization Buffer (eBioscience) and incubated in this buffer for 45 minutes at 4°C. Next, samples were washed twice in Permeabilization Buffer (eBioscience). The samples were then blocked for 15 minutes at 4°C in 100 μl of mouse and rat sera diluted in Permeabilization Buffer, washed once with Permeabilization buffer, and incubated for 45 minutes at 4°C in a 100 μl cocktail of intracellular antibodies (Table S2) diluted in Permeabilization Buffer. The cells were then washed with CyFACS and stained for 20 minutes at room temperature with 250 nM of Cell-ID^™^ Intercalator-IR (Fluidigm). Finally, the cells were washed twice with CyFACS buffer, once with MaxPar® cell staining buffer (Fluidigm), and once with Cell acquisition solution (CAS, Fluidigm), and then resuspended in EQ™ Four Element Calibration Beads (Fluidigm) diluted in CAS. Sample concentration was adjusted to acquire at a rate of 200 - 350 events/sec using a wide-bore (WB) injector on a CyTOF2 instrument (Fluidigm) at the UCSF Parnassus flow core facility.

### CyTOF data export and PP-SLIDE analysis

The CyTOF data were exported as FCS files, and samples were de-barcoded according to manufacturer’s instructions (Fluidigm). Events corresponding to antigen-specific cells were identified by gating on live, singlet intact CD3+CD19-CD4+ or CD8+ T cells expressing IFNγ, and exported as the population of antigen-specific cells. Data export was conducted using FlowJo (BD Biosciences) and Cytobank software. t-SNE analyses were performed using the Cytobank software with default settings. All cellular markers not used in the upstream gating strategy were included in generating the t-SNE plots. Non-cellular markers (e.g., live/dead stain) and the cytokines IFNγ, IL4, and IL17 were excluded for the generation of t-SNE plots. Dot plots were generated using both Cytobank and FlowJo.

The clustering method density-based clustering aided by support vector machine (DensVM) (Becher et al., 2014) was implemented in R to cluster, for each of the CD4+ or CD8+ T cell populations, total atlas cells, SARS-CoV-2-specific cells, or CMV-specific cells. DensVM builds on the method used by ACCENSE (Shekhar et al., 2014) for categorizing subpopulations, and calculates density by transforming the t-SNE data using the Gaussian kernel transformation.

PP-SLIDE analysis to define the predicted original states of antigen-specific IFNγ-producing cells followed our previously described method (Cavrois et al., 2017; Ma et al., 2020a). Briefly, using a custom script in R, each responding IFNγ+ cell from the stimulated sample was matched against every cell in the baseline unstimulated atlas phenotyped immediately after sample procurement, and then k-nearest neighbor (kNN) calculations were used to identify the phenotypically most similar cell in the atlas. The identified atlas cells harbor the predicted phenotypes of the original antigen-specific cells prior to IFNγ induction.

### CFSE Proliferation Assay

Cryopreserved PBMCs from two convalescent donors (4102 and 4106) were resuspended at 2×10^7^ cells/ml in PBS containing 0.1% FBS. The cells were then loaded for 10 minutes in the dark with 1 μM of CFSE (eBioscience). Labeling was stopped by adding 5 volumes of RP10. The cells were then incubated at 4°C for an additional 5 minutes. After 3 washes in RP10, the cells were cultured for 5 days in RP10 in the absence or presence of 10 ng/ml human IL-7 (R&D System). Cells were then stimulated for 6 hours with 0.5 μg/ml anti-CD49d clone L25 and 0.5 μg/ml anti-CD28 clone L293 (BD Biosciences), in the absence or presence of 0.5 μm PepMix™ SARS-CoV-2 Peptide (Spike Glycoprotein). Stimulation was conducted in the presence of Brefeldin A (3 μg/ml, eBioscience) to enable intracellular detection of induced IFNγ. Following the stimulation, cells were stained with APC/Cyanine7 anti-human CD3 (Clone SK7, Biolegend), PE/Dazzle^™^ 594 anti-human CD4 (Clone RPA-T4, Biolegend), Alexa Fluor 700 anti-human CD8 (Clone SK1, Biolegend), PE anti-IFNγ (Clone B27, BD Biosciences), and Zombie Violet Pacific Blue (Biolegend) as a Live/Dead discriminator. Stained cells were fixed and analyzed by FACS on an LSRII (BD Biosciences). Flowjo (BD Biosciences) was used for analysis. Live, singlet CD3+CD4+CD8-cells were assessed for proliferation by monitoring the loss of CFSE signal, and for the presence of SARS-CoV-2-specific cells by induction of IFNγ.

### Raw Data Availability

Raw CyTOF datasets used in this study are publicly available through the following link: https://datadryad.org/stash/landing/show?id=doi%3A10.7272%2FQ6RJ4GPP.

## Graphical Abstract

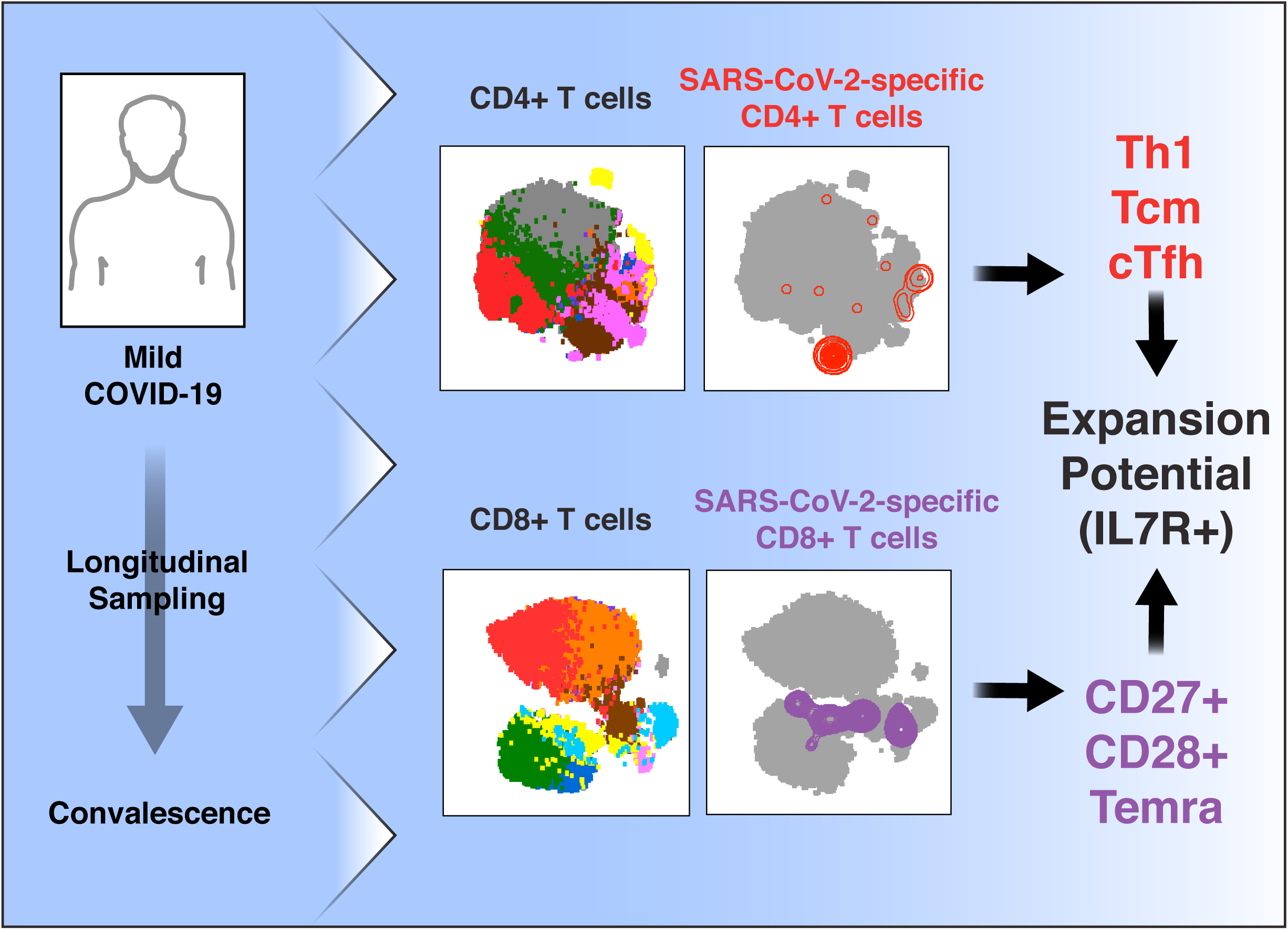

